# Neuromedin U Receptor 2 orchestrates sleep continuity and circadian entrainment

**DOI:** 10.1101/2025.08.30.673157

**Authors:** Konstantinos Kompotis, Sabrina Androvic, Miho Sato, Simon Gross, Marie Bainier, Rosa María Rodríguez Sarmiento, Bianca D Van Groen, Christophe Grundschober, Roger L Redondo, Steven A Brown

## Abstract

Neuromedin S signaling has recently been identified as a modulator of both sleep and circadian rhythms in the central nervous systems of mammals. Nevertheless, the involvement of its preferred receptor, NMUR2, in sleep and circadian regulation remains elusive. Here, employing its only selective antagonist R-PSOP, we demonstrate that NMUR2 function in the mouse brain is essential for physiological sleep architecture under baseline conditions, as well as for responses to homeostatic and circadian challenges. Using a combination of pharmacological, neuronal tagging, brain clearing and chemogenetic tools, we identified and functionally validated a group of PVH neurons as one of the targets mediating the effect of R-PSOP on sleep. These findings establish NMUR2 as a potential therapeutic target for the integral treatment of sleep and circadian neurological disorders.

**HIGHLIGHTS:** Selective NMUR2 antagonism via R-PSOP disrupts vigilance state continuity in a dose- and circadian phase-dependent manner

R-PSOP administration delays circadian phase entrainment and hinders sleep homeostasis recovery

PVH neurons are activated by R-PSOP and boost wake-to-NREM sleep transitions in the active period

## INTRODUCTION

The optimal coordination of mammalian processes and behaviors such as sleep, feeding, and temperature regulation throughout the 24-hour day relies heavily on neuropeptidergic signaling systems^1^. Amongst these systems, neuromedins occupy an increasingly central role as modulators of a plethora of physiological processes^2^. Their most studied family members, neuromedin U (NMU) and S (NMS), have been implicated in energy homeostasis^3^, stress responses^4^, circadian regulation^5^, as well as sleep and arousal from zebrafish to mice and rats^6–8^. While the role of NMU in feeding behavior and energy homeostasis has been well established, NMS has emerged as a central neuropeptide in circadian pacemaking^5^ and seasonal adaptation^9^, suggesting distinct functional roles for the two signaling pathways. In the context of circadian orchestration of sleep and arousal, the diurnal variation in NMS expression within the suprachiasmatic nucleus (SCN) of nocturnal rodents is particularly intriguing, with peak levels occurring during the end of the light phase^10^, and NMS signaling in hypothalamic circuits promoting arousal^11^. While both NMU and NMS selectively bind to specific G protein-coupled receptors (GPCRs), namely neuromedin U receptor 1 (NMUR1) and 2 (NMUR2), NMS exhibits a significantly higher affinity for NMUR2^12^, which is primarily expressed in the central nervous system and highly abundant in hypothalamic nuclei^13^. Interestingly, NMUR2 has been found to mainly couple to Gαq/11^14^, with some evidence suggesting its coupling to the inhibitory Gαi protein, at least in humans^15^.

The multifaceted role of neuromedin signaling in thermoregulation, energy balance and arousal, make NMUR2 ligands particularly interesting to integrally treat sleep and circadian disorders. The anatomical distribution of NMUR2 in hypothalamic circuits, including the SCN, positions it as a candidate for modulating sleep quantity and quality. Its role in the modulation of circadian rhythms and sleep has been investigated solely via the application of synthetic NMUR2 agonists, resulting in the enhancement of alertness and fragmentation of sleep^8^. We evaluated several tool compounds reported as NMUR2 antagonists including those described by Takeda (US 2006/0252797), Taisho (WO 2009/011336) and R-PSOP^16^. Among these, we confirmed that only R-PSOP was t acting as a selective small molecule NMUR2 antagonist. Conversely, the effects of R-PSOP, a selective small-molecule NMUR2 antagonist^16^, have not yet been studied in the context of sleep or circadian regulation. Despite its breakthrough discovery as the only selective NMUR2 antagonist and its use in a pain model administered intrathecally in rats, R-PSOP does not possess the necessary properties for its use in oral *in vivo* characterizations. Essential physicochemical and pharmacological properties needed for the therapeutic use of R-PSOP as a tool compound, such as permeability, brain penetration, bioavailability, route of administration and dosage for effective exposure in sleep models had to be addressed. Moreover, the precise role of NMUR2 in regulating circadian entrainment, sleep architecture and EEG has not yet been dissected. Similarly, the precise neural circuits through which NMUR2 exerts its effects have only recently started to be characterized.

The current study investigates the role of NMUR2 as a critical regulator of mammalian sleep-wake behavior, employing a combination of chrono-pharmacological interventions with R-PSOP, electroencephalography and the TRAP2 neuronal tagging system^17^, in conjunction with whole brain clearing and chemogenetics. We initially report the pharmacodynamic properties of R-PSOP in the brain and demonstrate its dose-dependent effect on locomotor activity (LMA) and sleep behavior. Antagonism of NMUR2 modulates sleep architecture in a bidirectional manner, depending on the circadian phase at which R-PSOP is administered. Beyond baseline sleep, NMUR2 antagonism is able to significantly alter the response to sleep homeostatic challenges and affects circadian entrainment in response to phase shifts. In search of the circuitry underlying the observed behaviors, we successfully identify neuronal ensembles activated by R-PSOP in the paraventricular hypothalamic nuclei (PVH), amongst other structures, also in a dose-dependent manner. Finally, chemogenetic reactivation of PVH neurons initially activated by R-PSOP application establishes a causal link between the PVH and specific aspects of sleep architecture and EEG activity altered by NMUR2 antagonism.

Our findings reveal a previously hinted, but thus far mechanistically elusive, role for NMUR2 in sleep behavior. The current study expands our understanding of the neural substrate controlling sleep and arousal, with important implications for understanding and treating systems modulating integral processes such as sleep.

## RESULTS

### Selectivity, pharmacokinetic and pharmacodynamic predictions of the NMUR2 antagonist R-PSOP

In terms of physicochemical properties, R-PSOP is a highly soluble compound (LYSA: 350 µg/mL); however, it exhibits very low permeability due to the presence of the urea linker **(Figure 1A**). In addition it has a moderate Pgp efflux ratio in rodents that limits the brain exposure for the compound (Pgp ER: 2.6). In fact it has been reported that R-PSOP has been used in a pain model administered intrathecally in rats^16^ due to the limited brain penetration of the compound. To address the permeability and the Pgp, analogs were synthesized, including modifications and alkylations of the urea linker to improve permeability. Despite these efforts, all modified analogs showed reduced or no potency at the NMUR2 receptor. On *in vitro* liver models (microsomes & hepatocytes) the compound showed low clearance (rate of elimination of the compound in the body). Nevertheless, we evaluated R-PSOP in a single-dose pharmacokinetic (SDPK) study in mice using different routes of dosage: intravenous (i.v.) (1mg/kg), oral (p.o)., and intraperitoneal (i.p.) (both 5mg/kg) (**Figure 1B**). We observed that *in vivo* clearance was higher than expected, likely due to extrahepatic clearance pathways and the compound’s low permeability in *in vitro* cellular assays, which affected the measured *in vitro* clearance On the PK *in vivo* study R-PSOP exhibits moderate to high clearance (41mL/min/Kg), moderate i.p. bioavailability (∼30%) and low oral bioavailability (2%) (**Figure 1B**). Although the compound has a short plasma half-life (T½ =1 hour), we thought that sufficient exposure can still be achieved within the first hour to observe a pharmacodynamic (PD) effect. The high clearance, the medium Pgp efflux ratio (AP-ER: 2.6) in rodents and the low permeability contribute to limited brain exposures determined by a free brain to plasma exposure ratio or Kp,uu of ∼15% (key properties in **Figure 1A(3)**). Considering the different bioavailabilities, the Kp,uu the fraction unbound in mice (fu: 36%) and the *in vitro* mouse NMUR2 binding (IC50= 197nM, N= 6) (**Figures 1A (2,3)** and **1B**), we estimated that in order to have a brain NMUR2 receptor occupancy above 80% between ∼0.5-1.2 hours post dosing a dose of more than 80mg/kg (e.g. 100mg/kg) with i.p. drug application will be needed. This was confirmed by a follow up mice PK with 100mg/kg i.p.dose using a solution prepared in 10% cyclodextrin (HP-beta-CD) in water with 0.7% sodium chloride as the vehicle (**Figure 1C,D**). In regard to the selected receptor occupancy, similar to other G-protein coupled receptors, we expected biological effects of NMUR2 antagonists to be observed between 70 to 90% receptor occupancy^18^.

**Figure 1.**
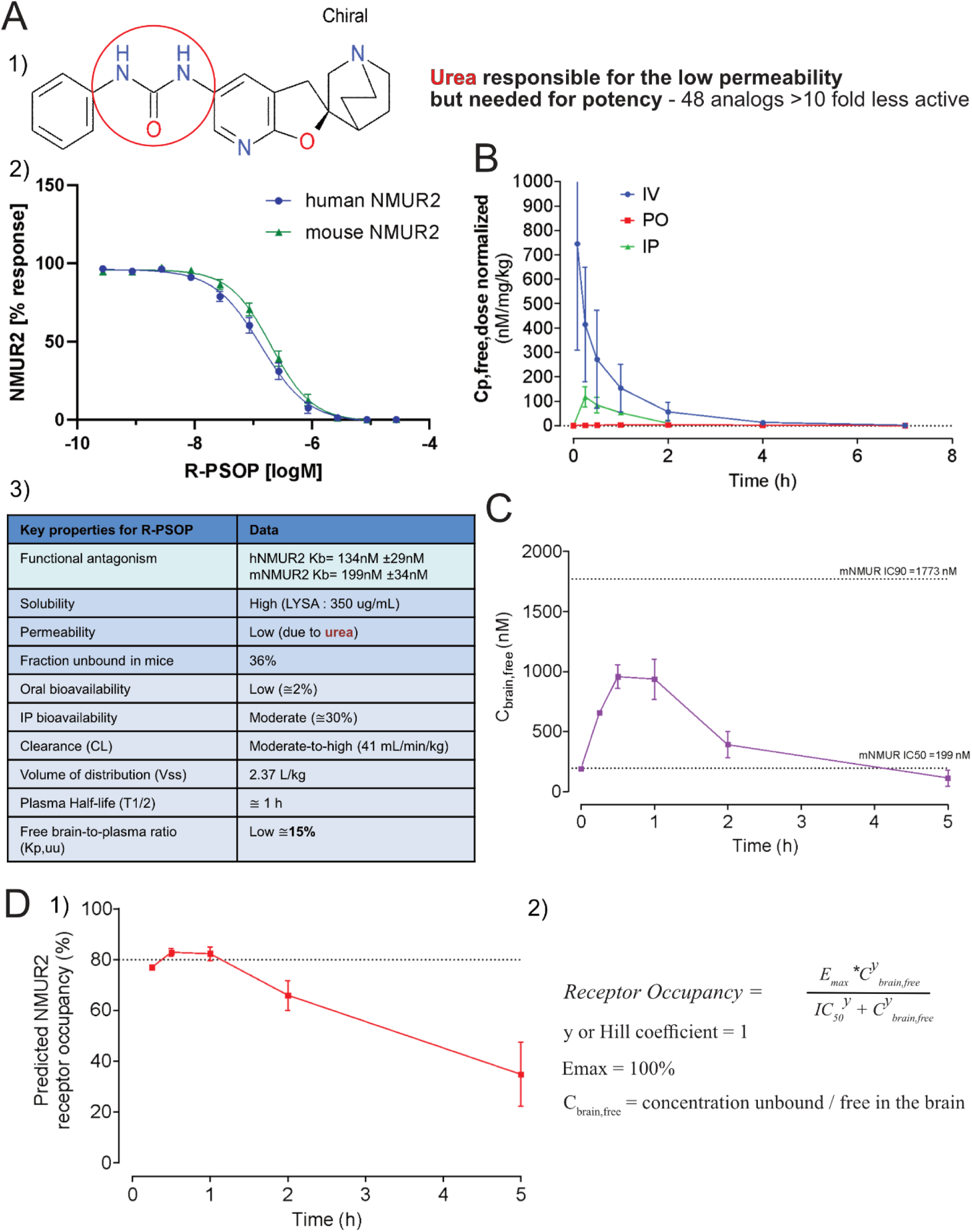
Profile for R-PSOP; measured in vitro, physicochemical and pharmacokinetic properties. (A) 1) Structure, key urea feature affecting permeability; 2) *In vitro* functional dose-response on human and mouse NMUR2 receptor; 3) *In vitro* data and key physicochemical & pharmacokinetic properties that affects *in vivo* profile. (B) Free plasma exposures with different routes of administration. (C) Pharmacokinetics and free brain exposure at 100 mg/kg i.p administration. (D) 1) Predicted receptor occupancy (∼80% between 0.5-1.2 hrs after administration of 100 mg/kg i.p in mice); 2) Calculation for the receptor occupancy. See also **Table S1**.

In terms of *in vitro* binding affinity R-PSOP has IC50: 131 nM at the human NMUR2 receptor and shows selectivity at the NMUR1 receptor (IC50: 27 μM). Broader selectivity of R-PSOP at 10 μM was assessed on a panel of 31 human and 5 rat receptors as well as in 13 human and 1 bovine enzyme assays (**Supplemental Table S1**). At this concentration, R-PSOP strongly inhibited radioligand binding to the human 5-HT3 by 94%, Muscarinic M1 by 59% and opioid Kappa receptor by 58%. This confirms the results by Liu and colleagues^16^ obtained on a panel of 104 human and rat receptors, transporters and enzymes. R-PSOP has been reported to bind to the 5-HT3 receptor (Ki= 220nM) which resulted in a receptor activation with EC50 of 6.8 μM in electrophysiological assays. It also binds to the Nicotinic alpha7 receptor (Ki= 460 nM) with an EC50 of 9.5 μM in electrophysiology (not tested in our study). Importantly, we did not reach μM concentrations of R-PSOP in our experiments. Additionally, R-PSOP showed no activity at 10 μM across all other tested receptors, transporters and enzymes. Overall, these findings confirm that R-PSOP can be used as a selective NMUR2 antagonist, as reported in the literature.

### R-PSOP application during the active period reduces locomotor activity and increases NREMS in a dose-dependent manner

NMUR2 agonists have been shown to increase activity and reduce sleep in rats, similar to the effect of NMS dosage. To investigate the impact of NMUR2 antagonism on sleep-wake behavior, we injected mice with different doses of R-PSOP, namely 10, 50 and 100mg/kg, two hours after dark onset (Zeitgeber Time 14; ZT14), and monitored their LMA and sleep behavior. Quantitative analysis revealed that R-PSOP at 50 and 100mg/kg significantly reduced LMA to less than 33% of the values observed after vehicle injection across the first 4 hours following compound application while the 10mg/kg dose had no apparent effect (**Figure 2A**, left upper panel). Accordingly, time spent in wakefulness decreased by 47.6 ± 9.9 and 36.9 ± 7.5 minutes as compared to vehicle, with no significant changes occurring under any other condition (**Figure 2A**, left bottom panel, and **Figure 2B**). Time spent in NREMS was increased by 46.6 ± 9.1 minutes and 37.3 ± 7.1 minutes, respectively, suggesting a somnogenic effect of R-PSOP (**Figure 2A**, right upper panel, and **Figure 2B)**. No differences in REMS amounts were observed in the hours following the injection; however, mice administered specifically with 100mg/kg R-PSOP exhibited increased REMS amounts during ZT18-20 compared to vehicle (**Figure 2A**, right bottom panel).

**Figure 2.**
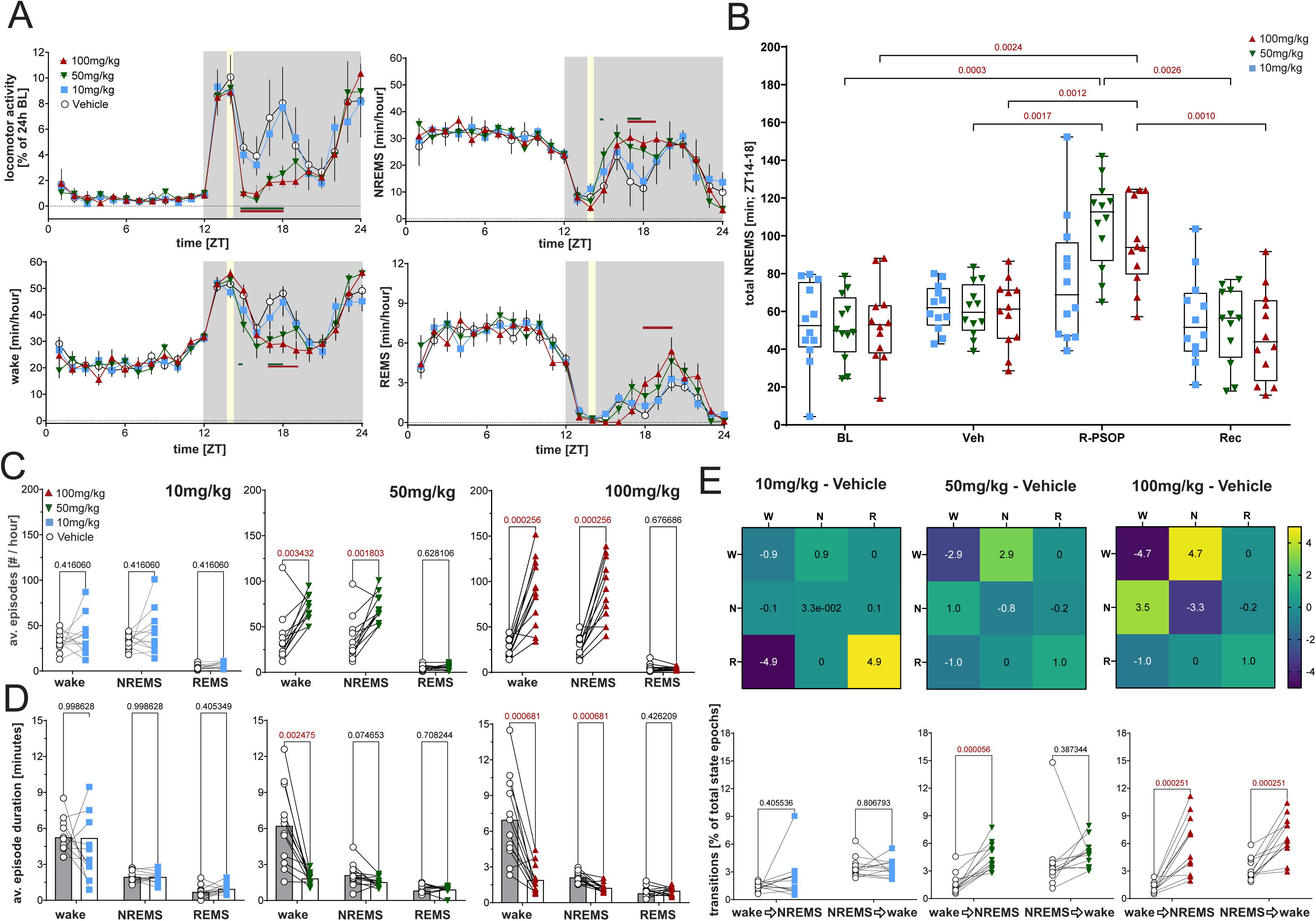
R-PSOP affects locomotor activity and sleep architecture in a dose-dependent manner. (A) Time course of locomotor activity (upper left), wakefulness (bottom left), NREM sleep (upper right), REM sleep (bottom right), and wakefulness over 24 hours in mice treated at ZT14 with vehicle (empty circles), 10mg/kg (blue squares), 50mg/kg (green inverted triangles), and 100mg/kg (red ascending triangles) of R-PSOP. Data points represent mean ± SEM. Gray shaded areas indicate dark phase; yellow vertical lines indicate time of intraperitoneal injection. Horizontal bars indicate significant differences between vehicle and R-PSOP-treated groups (green for 50mg/kg, red for 100mg/kg; 2-rANOVA, post-hoc Tukey tests, p < 0.05; n = 12). (B) Total NREMS duration between ZT14-18 in baseline (BL), vehicle (Veh) or R-PSOP injection, and recovery (Rec) day for the three different doses. Individual data points are shown as 10mg/kg (blue squares), 50mg/kg (green inverted triangles), and 100mg/kg (red upward triangles). Box plots show median and interquartile range; whiskers represent 10th and 90th percentiles. Statistically significant comparisons are shown above brackets in red (2-rANOVA, post-hoc Tukey tests, p < 0.05; n = 12). (C) Average episode number of wake, NREMS, and REMS in the 4 hours following administration of vehicle or R-PSOP. Individual data points are shown as empty circles (Vehicle, all panels), blue squares (10mg/kg, left panel), green inverted triangles (50mg/kg, mid panel), and red upward triangles (100mg/kg, right panel). Individual paired data points are connected by lines. Bar graphs represent mean ± SEM. Statistically significant differences are indicated above brackets in red (paired t-tests, p<0.05; n = 12). (D) Average episode duration (minutes) of wake, NREMS, and REMS in the 4 hours following administration of vehicle or R-PSOP. Individual data points are shown as empty circles (Vehicle, all panels), blue squares (10mg/kg, left panel), green inverted triangles (50mg/kg, mid panel), and red ascending triangles (100mg/kg, right panel). Individual paired data points are connected by lines. Bar graphs represent mean ± SEM. Statistically significant differences are indicated above brackets in red (paired t-tests, p<0.05; n = 12). (E) Heatmap showing changes in wake, NREMS and REMS transition probabilities (% of total state epochs) between vehicle and R-PSOP at 10mg/kg (left), 50mg/kg (middle), and 100mg/kg (right) for the 4 hours after injection. For each transition, the starting epoch state is indicated on the left, while the ending epoch state is indicated on top. Color scale indicates magnitude and direction of changes in transition probability (blue: decrease; yellow: increase). Bottom panels show individual animal transitions from wake-to-NREMS and NREMS-to-wake states for each respective heatmap. Corresponding p-values for significant differences are shown above brackets in red (paired t-tests, p<0.05; n = 12).See also **Figure S1**.

To unveil the source of the observed differences in wake and sleep amounts we further investigated sleep architecture in the hours following R-PSOP injection. The lowest dose had no significant effect on average state episode duration or number (**Figures 2C** and **D**, left panels). At 50mg/kg, R-PSOP doubled average wake and NREM episodes (+ 34.3 ± 8.3 and + 33.8 ± 7.1 from vehicle, respectively; **Figure 2C**, middle panel) and significantly reduced wake episode duration by 5.1 ± 0.9 minutes (**Figure 2D**, middle panel) as compared to vehicle treatment, indicating fragmented wakefulness. Notably, the maximum dose of R-PSOP increased average wake and NREM episode numbers by more than threefold (+ 62.0 ± 10.5 and + 61.4 ± 10.2 from vehicle, respectively; **Figure 2C**, right panel), while significantly reducing average wake and NREM episode duration (**Figure 2D**, right panel), reflecting a dose-dependent disruption in the continuity of both states. State transition analysis further confirmed the effect of NMUR2 antagonism on state maintenance, with 50mg/kg increasing the percentage of wake-to-NREMS transitions (+ 2.9%; **Figure 2E**, middle panel, and **Figures S2A** and **B**). At 100mg/kg both wake-to-NREMS and NREMS-to-wake transitions increased (+ 4.7% and +3.5%, respectively; **Figure 2E**, right panel, and **Figures S2A** and **B**) compared to vehicle.

Spectral analysis demonstrated that R-PSOP dose-dependently affected EEG composition (**Figures 3A and B**). More specifically, a characteristic EEG signature at 3.0 Hz during both wakefulness and NREMS emerged at the two higher doses (**Figure 3A**, middle and right panels). During NREMS in the two hours following injection of 50 mg/kg (middle panels) and 100 mg/kg (right panels) R-PSOP, EEG spectral power decreased mainly in the 1.5- to 2.5-Hz, 3.5- to 6.0-Hz, and 9- to 17.0-Hz bands (**Figure 3B**) compared to vehicle, while the lowest dose (10 mg/kg, left panels) produced no changes in EEG profiles. Since the NREMS EEG fast delta power (d2; 2.5–4.5Hz) is a proxy of homeostatic sleep pressure correlating well with the dynamics of sleep intensity^19^, we next followed its dynamics during the injection day. We observed that 50 and 100mg/kg of R-PSOP significantly reduced d2 power (−32% and −46.8% of vehicle, respectively) in the hour following compound administration, with the suppression lasting for one hour longer in the 100mg/kg-treated group (**Figure 3C**). To further assess the impact of NMUR2 antagonism on sleep depth, we employed a workflow for unsupervised clustering of sleep states (WUCSS)^20^, which can discriminate between deep and light sleep in mice. First, analysis of NREMS EEG spindle dynamics across the injection day confirmed that sleep quality was reduced in the first two hours after injection of the two higher R-PSOP doses, followed by a short increase in spindle density in the fourth hour (**Figure 3D**). Subsequent analysis showed that the gained NREMS time in the first two hours after injection was mostly light NREMS (**Figure 3E**, left panel), also supported by the observed reduction in d2 power (**Figure 3C**), while the last two hours contained more profound sleep (**Figure 3E**, right panel).

**Figure 3.**
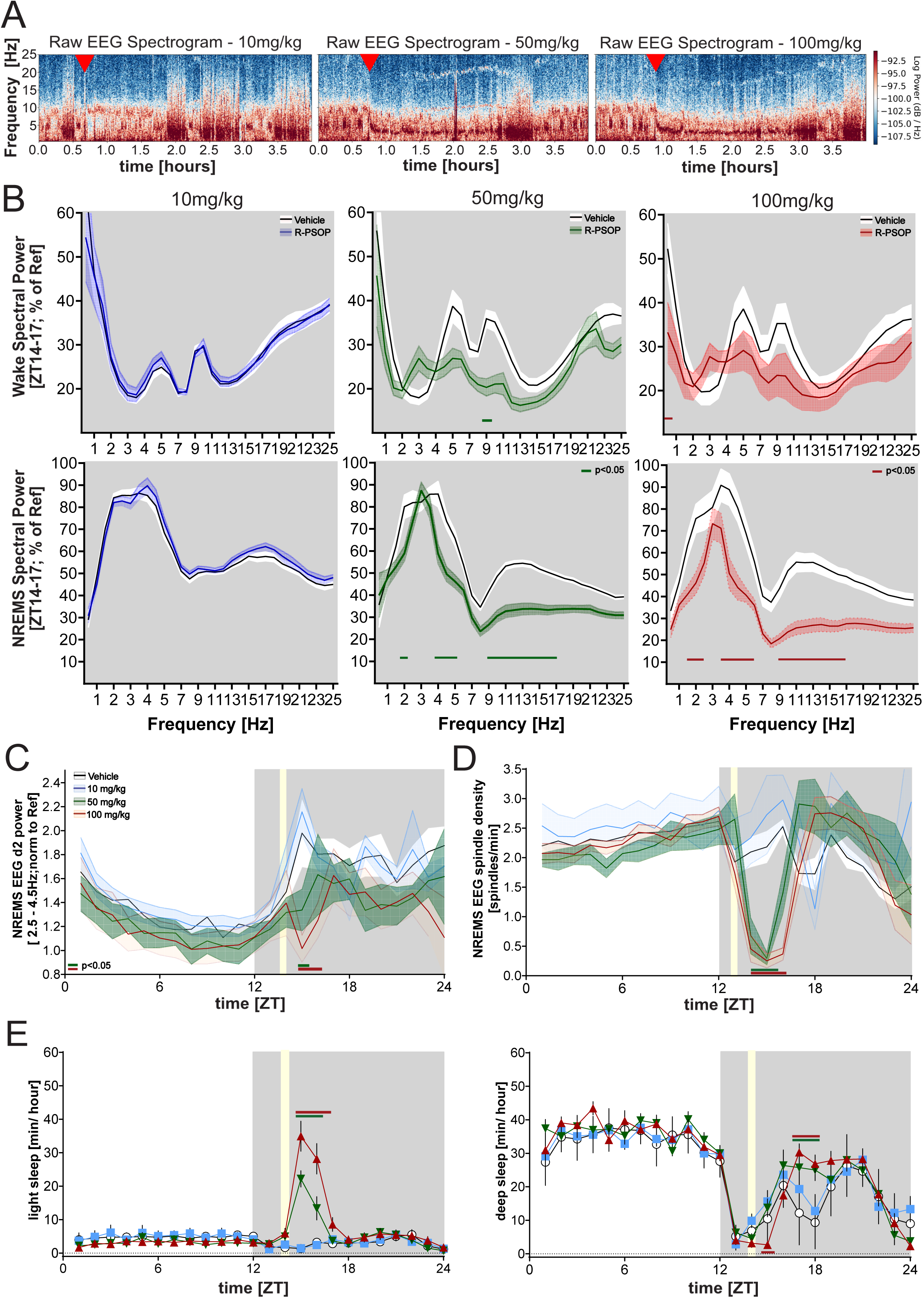
Dose-dependent effects of R-PSOP on sleep EEG and quality. (A) Representative raw EEG spectrograms depicting the hour prior to and the first 3 hours following administration of R-PSOP at 10mg/kg (left panel), 50mg/kg (mid panel), and 100mg/kg (right panel). Red arrowheads indicate time of injection. Color scale represents log power. (B) Wake (top panels) and NREMS EEG spectral power (ZT14-16; % of BL reference; see Methods) in the ZT14-17 period, following administration of 10mg/kg (blue line, left panels), 50mg/kg (green line, middle panels), and 100mg/kg (red line, right panels) R-PSOP compared to vehicle (black lines). Shaded areas represent SEM. Horizontal colored lines indicate frequencies with significant differences between R-PSOP and vehicle treatments (Mixed effects model; post-hoc Bonferroni, p<0.05; n = 12). (C) Hourly dynamics of NREMS delta2 power (2.5-4.5Hz; normalized to EEG d2 of BL ZT8-12; also see Methods) across the day of vehicle (black) or R-PSOP at 10mg/kg (blue line), 50mg/kg (green line), or 100mg/kg (red line) administration. Shaded areas represent SEM. Gray shaded areas indicate the dark period, while the yellow shaded area represents the injection window at ZT14. Statistically significant differences between the vehicle and the 50mg/kg or 100mg/kg-treated group are indicated by the horizontal green and red lines, respectively (2-rANOVA; post-hoc Tukey, p<0.05; n=12). (D) Hourly spindle density during NREMS across the day of vehicle (black) or R-PSOP at 10mg/kg (blue line), 50mg/kg (green line), or 100mg/kg (red line) administration. Data are shown as mean ± SEM. Gray shaded area represents the injection window at ZT14. Statistically significant differences between the vehicle and the 50mg/kg or 100mg/kg -treated group are indicated by the horizontal green and red lines, respectively (2-rANOVA; post-hoc Tukey, p<0.05; n=12). (E) Hourly dynamics of deep sleep time (left panel) following administration of vehicle (black) or R-PSOP at 10mg/kg (blue line), 50mg/kg (green line), or 100mg/kg (red line). On the right, hourly dynamics of light sleep time under the same conditions. Data are presented as mean ± SEM. Gray shaded area represents the injection window at ZT14. Statistically significant differences between the vehicle and the 50mg/kg or 100mg/kg -treated group are indicated by the horizontal green and red lines, respectively (2-rANOVA; post-hoc Tukey, p<0.05; n=12). See also **Figure S2A-D**.

Together, our findings suggest that NMUR2 antagonism during the active period affects both the quantity and quality of wakefulness and sleep in a dose-dependent manner, mainly boosting light sleep in mice by disrupting wake continuity. Since 100mg/kg of R-PSOP maximized the observed effect on behavior in both intensity and duration, we selected the maximum dose to characterize its role in the following experiments.

### Antagonism of NMUR2 during the resting period delays entrainment to circadian phase advance

Previous research has established a circadian regulatory function for NMS, the principal ligand of NMUR2, within the SCN^5,9^. We first set out to validate recent observations that SCN NMS neurons are necessary in arousal regulation^7^, and assess their involvement in our observed effects of NMUR2 antagonism on sleep and LMA. In our hands, sleep and wake behavior of mice with silenced SCN NMS neurons did not differ following administration of vehicle or 100mg/kg R-PSOP at ZT14, with the exception of a significant LMA reduction (**Figure S2A-D**), supporting the idea that physiological SCN NMS-NMUR2 signaling is vital for the optimal timing of sleep and wake regulation. To clearly elucidate the role of NMUR2 in the regulation of circadian behaviors, we next examined entrainment dynamics in a different cohort, receiving either vehicle or 100mg/kg R-PSOP prior to a 6-hour phase advance in the light-dark cycle (**Figure 4A**). Comparison of average actograms across a week after injection (**Figure 4B**) revealed entrainment differences between treatment groups. While vehicle-treated mice (blue-shaded plot) rapidly adapted their activity onset to the new schedule, R-PSOP-treated mice (pink-shaded plot) exhibited significantly delayed entrainment to the advanced dark onset (**Figure 4C**), taking almost a day longer than the vehicle-treated group to reach 50% of phase shift (3.5 vs 2.75, respectively; **Figure 4D**). Further investigation of the acute effect of R-PSOP dosage (**Figure 4E**) suggested that this delay in the R-PSOP-treated group might result from an LMA reduction during the first 2 hours following injection, although comparison of LMA for each 30-minute-bout of the first day did not exhibit any statistically significant differences between the two groups. These results indicate that physiological NMUR2 signaling plays a critical role in facilitating rapid entrainment to phase advances, and support its involvement in the molecular machinery that synchronizes internal circadian rhythms to environmental light cues.

**Figure 4.**
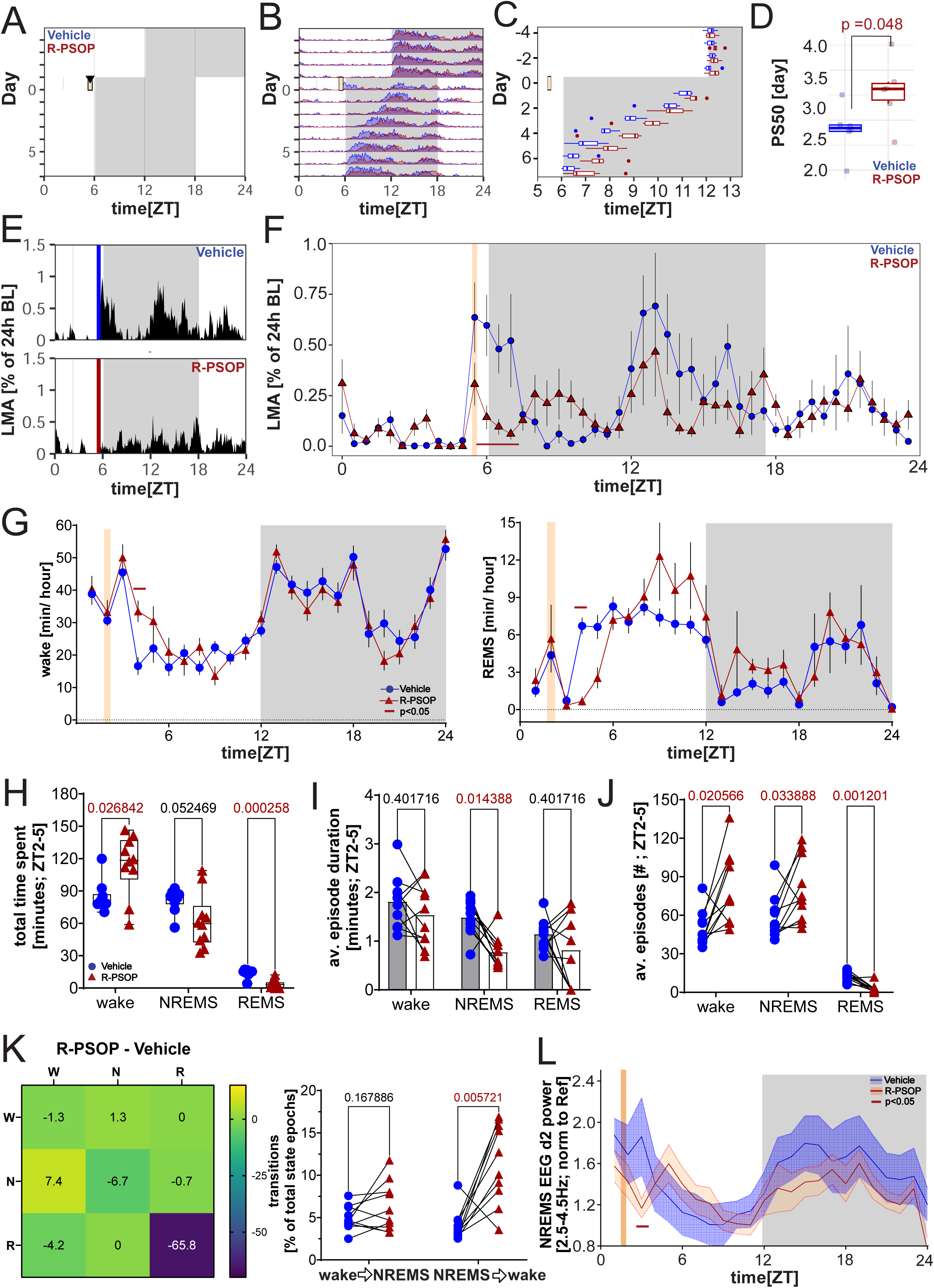
Effects of NMUR2 antagonism during the resting period on circadian entrainment and sleep-wake behavior. A) Schematic actogram showing the experimental design across multiple days. Yellow and gray background indicate light and dark periods, respectively. Black arrowheads indicate the day and time of treatment. Following 4 days of baseline recordings under 12:12 LD, light offset was advanced for 6 hours. Animals were treated with either vehicle (n=6) or 100mg/kg R-PSOP (n=8) at ZT6, prior to lights off. B) Day-by-day activity plots comparing entrainment to a 6-hour advance of light onset after treatment with vehicle (blue line) and R-PSOP (pink line) across the 24-hour day. Arrowhead indicates the day of administration. C) Analysis of activity onset times across days following vehicle (blue) or R-PSOP (pink) treatment. Each point represents an individual animal. Box plots indicate median and interquartile range. Arrowhead indicates day of treatment. D) Phase shift magnitude (PS50; time to reach 50% phase shift) measured in days following vehicle (blue) and R-PSOP (pink) administration. Box plots show median, 25th-75th percentiles, and min-max values. Individual data points are shown. Statistical significance indicated above brackets in red (t-test, P<0.05; n_vehicle_=6, n_R-PSOP_=8). E) Average locomotor activity (LMA; counts per minute) dynamics over the first 24 hours following vehicle (top, blue arrowhead) or R-PSOP (bottom, pink arrowhead) administration. F) Detailed analysis of locomotor activity counts per 30-minute bin across the 24-hour day. Individual data points and averages are shown for the vehicle (blue dots and boxes) and R-PSOP (pink dots and boxes). Black arrowhead indicates time of administration. No statistically significant changes were observed (2-ANOVA; post-hoc Bonferroni, p<0.05; n_vehicle_=6, n_R-PSOP_=8). G) Hourly dynamics distribution of wake (left) and REMS (right) following vehicle (blue circles) or R-PSOP (red triangles) administration at ZT2. Gray shaded areas indicate the dark period, while the yellow shaded area represents the time of injection of either vehicle or 100mg/kg R-PSOP. Statistically significant differences are highlighted by the dark red horizontal line (2-rANOVA; post-hoc Bonferroni, p <0.05; n = 10/group). H) Total time spent (in minutes) in wake, NREMS, and REMS within the first 3 hours following injection. Box plots show median, 25th-75th percentiles, and min-max values. Individual data points are shown for each animal in the vehicle (blue circles) or 100mg/kg R-PSOP (red triangles) -treated group. Statistically significant differences between the groups are indicated above the brackets in red (paired t-test, p<0.05; n=10/group). I) Average episode duration (in minutes) for wake, NREMS, and REMS during ZT2-5 following vehicle (blue circles) or R-PSOP (red triangles) administration. Individual paired data points are connected by lines. Statistically significant differences between the groups are indicated above the brackets in red (paired t-test, p<0.05; n=10/group). J) Average number of episodes for wake, NREMS, and REMS during ZT2-5 following vehicle (blue circles) or R-PSOP (red triangles) administration. Individual paired data points are connected by lines. Statistically significant differences between the groups are indicated above the brackets in red (paired t-test, p<0.05;n=10/group). K) Heatmap summarizing changes in wake, NREMS and REMS transition probabilities (% of total state epochs) between vehicle and R-PSOP, across the 3 hours after injection (left panel). For each transition, the starting epoch state is indicated on the left, while the ending epoch state is indicated on top. Color scale indicates magnitude and direction of changes in transition probability (blue: decrease; yellow: increase). On the right panel, individual animal transitions from the respective wake-to-NREMS and NREMS-to-wake states are shown for the vehicle (blue circles) or R-PSOP (red triangles) treated group. Individual paired data points are connected by lines. Statistical significance is indicated in red above the bracket (paired t-test, p<0.05; n = 10/group). L) NREMS EEG fast delta (δ2) power (2.5-4.5Hz; normalized to d2 during BL ZT8-12; also see Methods) throughout the injection day. Blue- and red-shaded lines represent vehicle and R-PSOP-treated groups, respectively. Gray shaded areas indicate dark phases, while the yellow shaded area indicates the time of vehicle or R-PSOP administration. Statistically significant observations are shown by the dark red horizontal line (Mixed effects model for ZT0-12; post-hoc Bonferroni, p<0.05; n = 8/group). See also **Figure S2E-G**.

### R-PSOP application during the resting period reduces total time spent in NREMS

Our findings on the NMUR2 involvement in circadian regulation, as well as previous observations of NMS dosage differentially impacting circadian rhythms in rats depending on the time of injection^10^, suggest a potentially reciprocal interaction between circadian rhythms and NMUR2 function. We thus hypothesized that NMUR2 antagonism produces distinct effects depending on the time of administration. Indeed, application of 100mg/kg R-PSOP two hours after light onset (ZT2) increased time spent awake (**Figure 4G**, left panel) mainly at the expense of REMS (**Figure 4G**, right panel and **Figure S2E**) during the subsequent three hours (+30.9 ± 10.9 and −11.4 ± 1.4, respectively; **Figure 4H**). Across the same period, we observed a decrease in the average duration of NREMS episodes (−0.7 ± 0.19 min), while the average number of both wake and NREMS episodes increased (+32.0 ± 10.0 and +22.0 ± 9.0, respectively), and REMS episodes decreased (−9.0 ± 2.0; **Figure 4I**) as compared to vehicle. Transitions between vigilance states also differed, with a significant surge observed mainly in NREMS-to-wake transitions compared to vehicle (from 4.0 to 11.4 %; **Figure 4K**). Finally, R-PSOP dosage reduced NREMS EEG d2 power compared to vehicle in the first hour following injection (−37% of vehicle; **Figure 4K**) and produced the characteristic 3.0-Hz EEG signature (**Figure S2F** and **S2G,** left panel), similar to effects observed during the active period (ZT14). These findings suggest that although R-PSOP affects sleep-wake stability and NREMS EEG in a manner primarily independent of circadian influence, the final impact of NMUR2 antagonism on sleep-wake behavior depends on the circadian context and particularly on the predominant vigilance state during the circadian phase of administration.

### R-PSOP alters responses to homeostatic sleep challenges

Due to the observed wake-promoting effect of NMUR2 antagonism in the early light period, we set out to investigate whether R-PSOP would favor wakefulness even under increased sleep pressure conditions during the resting period. To this end, we subjected animals to 4 hours of sleep deprivation (SD; ZT0-4) and administered either vehicle or R-PSOP (100mg/kg) close to the end of SD. Following the homeostatic challenge, LMA was reduced only within the first hour in mice treated with R-PSOP as compared to vehicle, while both hourly and total wakefulness and NREMS amounts for the subsequent three hours (ZT4-7) were similar to vehicle-treated controls (**Figure 5B**, top and mid panel, respectively, and **Figure 5C**, left panel). Conversely, hourly (**Figure 5B**, bottom panel) and total REMS amounts exhibited a significant reduction in the same period (−11.03 ± 1.8 min; **Figure 5C**, right panel). Further analysis of sleep architecture revealed a significant reduction in both average wake and NREMS episode duration (−1.49 ± 0.3 and −1.46 ± 0.2 minutes, respectively; **Figure 5D**) in the three hours following SD. For the same period, both average wake and NREMS episode number increased, with a simultaneous decrease in average REMS episode number (+48.8 ± 3.8, +52.2 ± 5.8 and −7.2 ± 1.5, respectively; **Figure 5E**). State transition analysis showed elevated wake-to-NREMS (+6.3%) and NREMS-to-wake (+4.3%) transition probabilities in the R-PSOP-treated compared to the vehicle-treated group (**Figures 5F**, right panel, and **S3A**). Suggesting a wake-disrupting effect, dosage of R-PSOP significantly reduced sleep-onset latency following SD by 13.9 ± 4.9 min (65%) compared to vehicle. Interestingly, a marked increase in NREMS EEG spectral power particularly in the 2.5-4.5 Hz range was observed in the first two hours of recovery (**Figures 5H** and **S3B**). Following up on the dynamics of this band across the baseline (BL) and SD day, we observed that NREMS EEG d2 power in the R-PSOP-treated group remained elevated in the hours after forced wakefulness and persisted until the end of the resting period (+33.5% of vehicle on average for ZT5-7; **Figure 5I**). Our observations suggest that NMUR2 antagonism results in impaired sleep consolidation despite homeostatic pressure and interferes with homeostatic sleep recovery mechanisms. They also demonstrate that the effect of R-PSOP on sleep architecture and EEG profiles persists towards the same direction regardless of circadian period or sleep-homeostatic pressure, while transition probabilities and vigilance state amounts differ depending on the circadian or homeostatic context.

**Figure 5.**
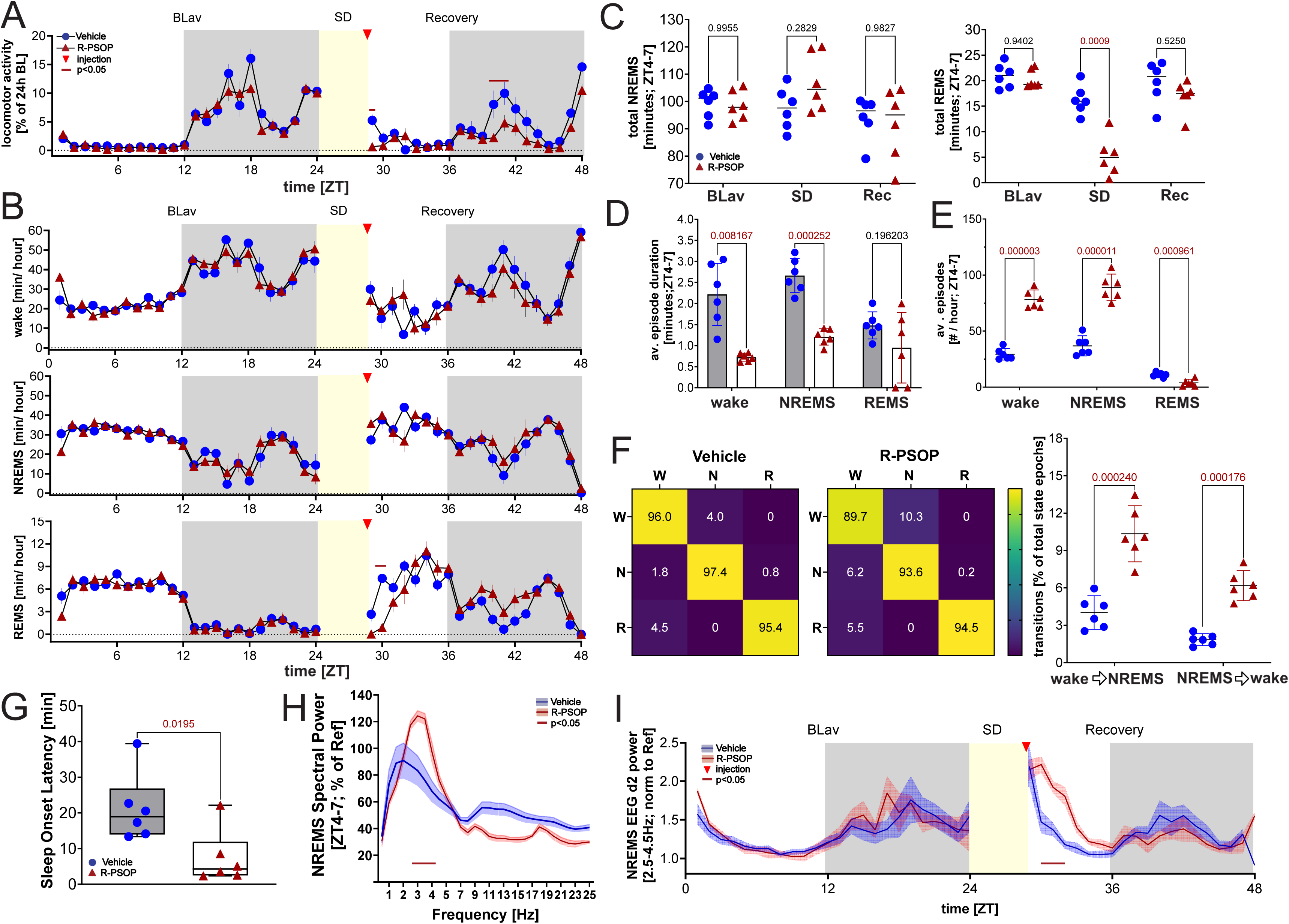
Administration of R-PSOP alters homeostatic response to forced wakefulness. (A) Hourly dynamics of locomotor activity across baseline days (BLav), 4-hour sleep deprivation (SD), and recovery (Rec) days. Blue circles and red triangles represent vehicle and R-PSOP treated groups, respectively. Gray shaded areas indicate dark phases, while the yellow shaded area represents SD duration (ZT0-4) and the red arrow indicates administration time of either vehicle or R-PSOP. Statistically significant observations are shown by the dark red horizontal line (2-ANOVA; post-hoc Tukey, p<0.05; n = 6/group). (B) Hourly dynamics of wake (top), NREMS (middle), and REMS (bottom) for the respective days as in (A). Blue circles and red triangles represent vehicle and R-PSOP -treated groups, respectively. Gray shaded areas indicate dark phases, the yellow shaded area represents the SD duration (ZT0-4) and the red arrow indicates administration time of either vehicle or R-PSOP. Statistically significant observations are shown by the dark red horizontal line (2-ANOVA; post-hoc Tukey, p <0.05; n = 6/group). (C) Total time spent in NREMS (left panel) and REMS (right panel) during ZT4-7 in the two baseline (BLav), sleep deprivation (SD) and recovery (Rec) days. Individual data points are shown for each animal in the vehicle (blue circles) or 100mg/kg R-PSOP (red triangles) group, while group means are shown as horizontal black lines. Statistically significant differences are indicated above brackets in red (t-tests, p<0.05; n = 6/group). (D) Average episode duration of wake, NREMS, and REMS in the 3 hours following SD and administration of vehicle or R-PSOP. Individual data points are shown for each animal in the vehicle (blue circles) or 100mg/kg R-PSOP (red triangles) group. Bar graphs represent mean ± SEM. Statistically significant differences are indicated above brackets in red (t-tests, p<0.05; n = 6/group). (E) Average episode number of wake, NREMS, and REMS in the 3 hours following SD and administration of vehicle or R-PSOP. Individual data points are shown for each animal in the vehicle (blue circles) or 100mg/kg R-PSOP (red triangles) group. Group means are shown as horizontal black lines. Statistically significant differences are indicated above brackets in red (t-tests, p<0.05; n = 6/group). (F) Heatmap representation of transition probabilities (%) between wake (W), NREMS (N), and REMS (R) for the vehicle (left) and R-PSOP (right) -treated groups for the 3 hours following the end of SD and injection. For each transition, the starting epoch state is indicated on the left, while the ending epoch state is indicated on top. Color scale indicates magnitude and direction of changes in transition probability (yellow: high; purple: low). On the right panel, individual animal transitions from wake-to-NREMS and NREMS-to-wake states for each respective heatmap are shown. Corresponding p-values for significant differences are shown above brackets in red (t-tests, p<0.05; n = 6/group). (G) Sleep onset latency (min) following SD and vehicle (blue circles) or R-PSOP (red triangles) treatment. Box plot shows median and interquartile range with individual data points. Statistical significance is indicated in red above the bracket (t-test, p<0.05; n = 6/group). (H) NREMS EEG spectral power (ZT4-6; % of baseline total EEG power; also see Methods) across frequencies (0.5-25 Hz). Blue shaded line represents vehicle-treated animals, red shaded line represents R-PSOP-treated animals. The red line indicates frequency bands with significant differences between groups (p<0.05; n = 6/group). (I) NREMS EEG fast delta (δ2) power (2.5-4.5Hz; normalized to EEG d2 of BL ZT8-12; also see Methods) throughout the recording days in (A). Blue- and red-shaded lines represent vehicle and R-PSOP-treated groups, respectively. Gray shaded areas indicate dark phases, while the yellow shaded area represents the SD duration (ZT0-4) and the red arrow indicates administration time of either vehicle or R-PSOP. Statistically significant observations are shown by the dark red horizontal line (Mixed effects model for ZT4-12; post-hoc Bonferroni, p<0.05; n = 6/group). See also **Figure S3**.

### Paraventricular hypothalamic neurons are activated following administration of R-PSOP in a dose-dependent manner

The effects of NMUR antagonism on sleep architecture observed in various circadian and sleep contexts in our hands suggested that there might be multiple components of the neuronal circuitry involved in its mediation, each one contributing to different aspects of sleep-wake behavior. We thus set out to identify neural substrates mediating the effects of NMUR2 antagonism in the mouse brain. Pursuing an initial agnostic approach, we aimed to examine neuronal activation following R-PSOP dosage in the whole brain. To this end, we first crossed fosTRAP2 (Fos2A-iCreER^17^) and tdTomato (Ai14) mice, which enabled us to label active neurons during R-PSOP injection at ZT14 (**Figure 6A**). Subsequently, we employed whole brain clearing (CUBIC-HV tissue clearing^21^; also see Methods), thereby rendering the brain transparent. Three-dimensional imaging (mesoSPIM^22^; also see Methods) of tdTomato-expressing “TRAPped” neurons in the transparent brain revealed substantial neuronal activation in the paraventricular hypothalamus (PVH), among other structures, with dense clustering of labeled cells visible across multiple anatomical views (**Figures 6B** and **S4A**). Conventional immunofluorescent imaging on brain sections from animals administered with vehicle (**Figure 6C**, left panels) or R-PSOP (100mg/kg; **Figure 6C**, right panels, and **S4B**) at ZT14 confirmed robust cFos expression particularly within the PVH of antagonist-treated mice, while some activation was also observed in vehicle-treated controls, as expected. Since the mouse PVH is known to be active during the active period and largely inactive during the subjective day (ZT0-12), we further investigated any dose-dependent effects of R-PSOP on PVH-neuronal activation by injecting animals at ZT2 with 100 or 50mg/kg R-PSOP (**Figure 6D**, top and middle panels, respectively), and comparing them to vehicle- and R-PSOP-10mg/kg-treated control mice (**Figure 6D**, bottom panel). Quantitative analysis (**Figure 6E**) demonstrated an increase in the absolute numbers of cFos-positive cells in the PVH of 100mg/kg-R-PSOP-treated mice compared to controls (258 vs 25 cells, respectively), with a potentially intermediate effect at 50mg/kg (119 vs 25 cells). These findings establish the PVH as a key neuroanatomical substrate responding to NMUR2 antagonism, with activation patterns that parallel at least some of the dose-dependent effects on sleep-wake behavior observed in our study.

**Figure 6.**
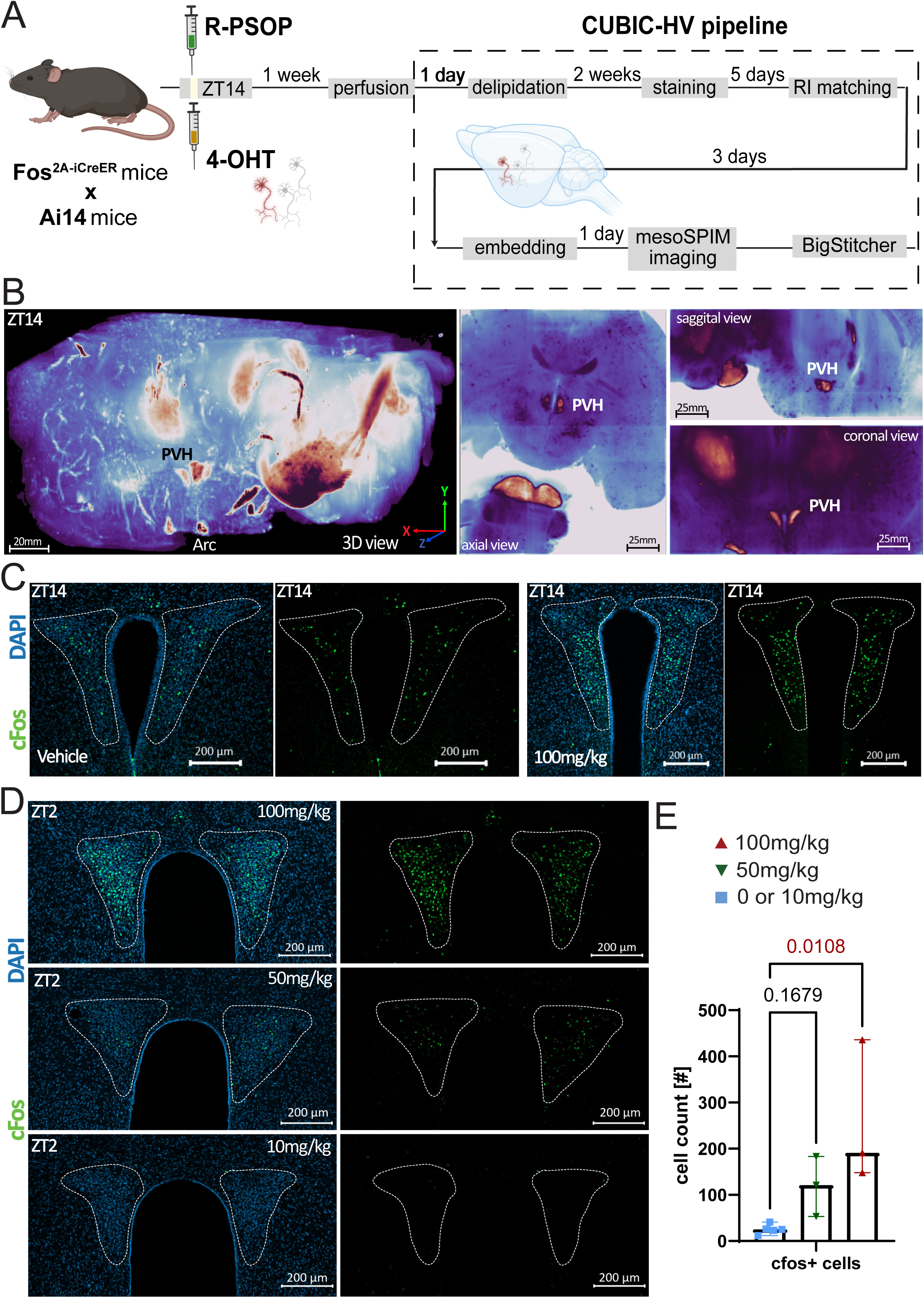
R-PSOP activates paraventricular hypothalamic neurons in a dose-dependent manner. (A) Experimental workflow for activity-dependent neuronal labeling in Fos2A-iCreER × Ai14 mice. Animals received intraperitoneally (i.p.) 4-OHT (20mg/kg) at the same time as R-PSOP (100mg/kg), allowing for permanent labeling of R-PSOP-activated neurons. After 1 week, brains were extracted, followed by delipidation, nucleic staining and RI matching using the CUBIC-HV tissue clearing pipeline, and subsequently imaged with mesoSPIM light-sheet microscopy for whole-brain visualization of activated neural populations (n=4). Files were pre-processed and stitched in BigStitcher (FIJI; also see Methods). (B) Three-dimensional visualization of R-PSOP-activated neurons throughout the brain in a representative sample. Multiple views [3-dimensional (3D), coronal, axial, and sagittal] show labeled neurons with particular enrichment in the paraventricular hypothalamus (PVH) and arcuate nucleus (ARC). Yellow/orange dots represent individual activated neurons; blue background shows brain autofluorescence for anatomical reference. Scale bar = 20 mm. Napari python package was used for visualization (also see Methods for specific parameters). (C) Representative immunofluorescent images of the paraventricular hypothalamus (PVH; white dashed lines) 1.5 hours after injection at ZT14 with either vehicle (two left panels) or R-PSOP (100mg/kg, two right panels). DAPI nuclear stain (blue) and cFos immunoreactivity (green) show increased neuronal activation in R-PSOP-treated brain. In each group, first the overlap of DAPI and cfos is presented, followed by the single cfos channel. Scale bar = 200 μm. (D) Representative immunofluorescent images of PVH (PVH; white dashed lines) show DAPI (blue) and cFos (green) immunofluorescence 1.5 hours following administration of 100mg/kg (top), 50mg/kg (middle), or 10mg/kg (bottom) R-PSOP at ZT2. In each group, first the overlap of DAPI and cfos is presented, followed by the single cfos channel. Scale bars = 200 μm. (E) Quantification of cFos-positive cells in the PVH across the different R-PSOP doses from (E). Individual cell counts are shown for each animal in the control blue squares; includes both 0mg/kg (vehicle) and 10mg/kg treatment], 50mg/kg (green inverted triangles) and 100mg/kg (red triangles) group. Bar graphs show mean ± SEM. Significant differences are shown above brackets in red (t-test, p<0.05, n_0/10mg/kg_ = 5, n_50mg/kg_ = 3, n_100mg/kg_ = 3). See also **Figure S4**.

### Reactivation of captured PVH neurons after R-PSOP dosage in the active period is sufficient to promote wake-to-NREMS transitions

The expression of NMUR2 in neuronal ensembles of the murine PVH has been previously demonstrated^9,23^. Following up on our observations of PVH-activated neurons post-R-PSOP administration, we hypothesized that **this** particular neuronal subset is functionally implicated, at least partially, in mediating the effect of NMUR2 antagonism. To test our hypothesis, we employed a combination of neuronal tagging and chemogenetics to first capture PVH neurons responsive to R-PSOP (100 mg/kg) administration at ZT14, similarly to our 3-D mapping experiment, and selectively reactivate them via clozapine-N-oxide (CNO; 4 mg/kg) application at the same timepoint a week later (**Figure 7A**). In mice with successful recombination and expression of hM3D(Gq)-mCherry (**Figure 7B**), this reactivation did not significantly reduce hourly LMA or alter state dynamics (**Figures 7C** and **D**), as previously observed under direct R-PSOP administration. However, it was sufficient to promote increased total NREMS amounts across the first 3 hours after **administration**, while total time spent awake decreased correspondingly (**Figure 7E**). Although sleep architecture was otherwise not significantly altered (**Figure S5A-C**), the probability of wake-to-NREMS transitions significantly increased after CNO application compared to saline (**Figure 7F**). Notably, the EEG signature observed following R-PSOP injection at ZT14 and ZT2 was absent after PVH reactivation (**Figure 7G**). Although wake and REMS EEG spectral power was identical between groups (**Figure 7H**, left panel, and **S5D**), NREMS EEG spectral power was significantly reduced in mice with reactivated PVH neurons for all bands above 2.0 Hz (**Figure 7H**, right panel), reminiscent of the reduction observed after administration of both 50 and 100 mg/kg R-PSOP at the same timepoint (**Figure 3B**, middle and right bottom panels). Lastly, NREMS EEG d2 power was reduced specifically in the first hour following PVH neuronal reactivation (−33% of vehicle; **Figure 7I**). Together, our observations establish that R-PSOP-responsive PVH neurons constitute a critical circuit node mediating the sleep-promoting effects of NMUR2 antagonism during the active period through increased transitions from wakefulness to NREM sleep. These findings also highlight that multiple structures are involved in mediating different aspects of its effect on sleep-wake regulation.

**Figure 7.**
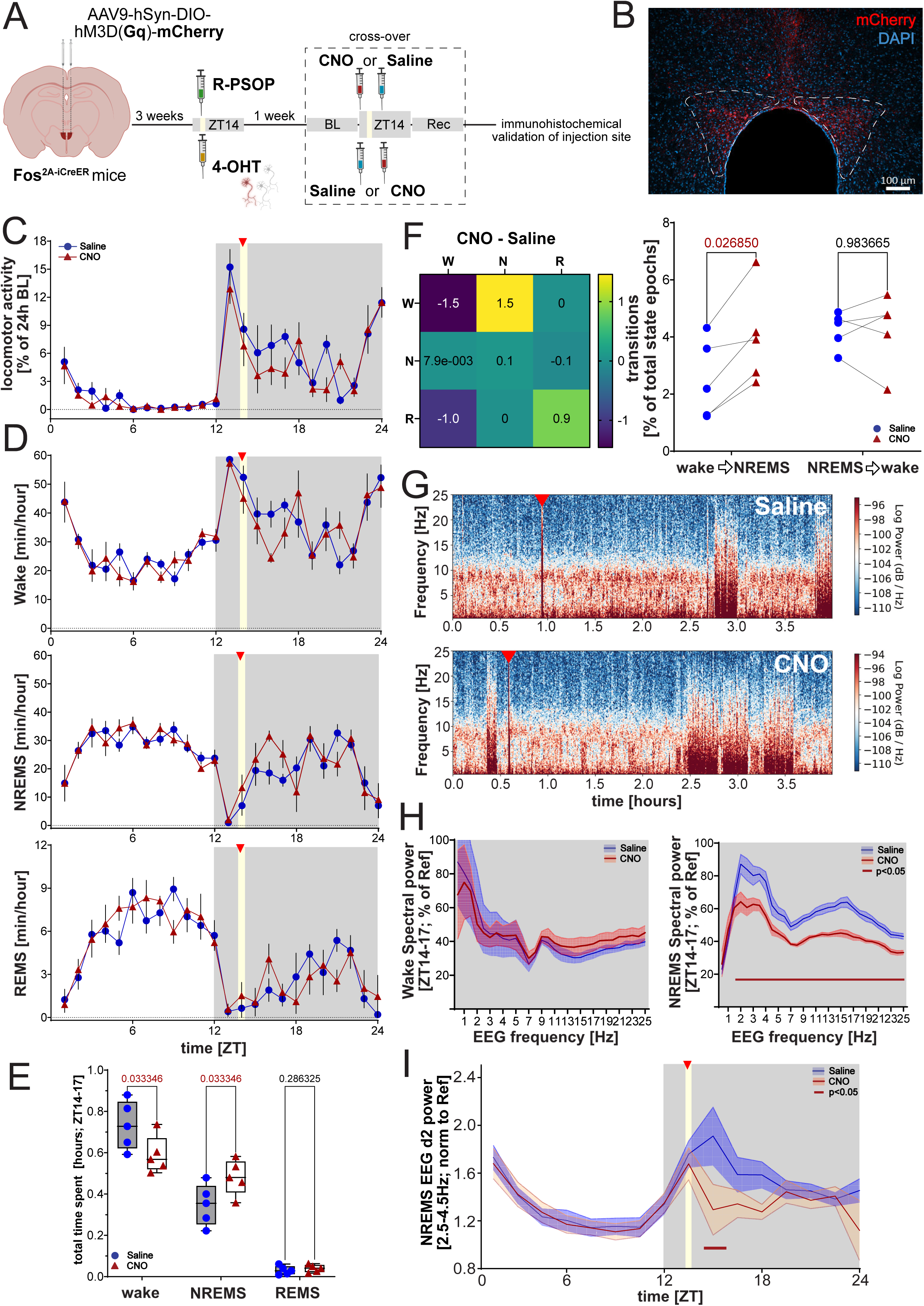
Chemogenetic reactivation of R-PSOP-responsive neurons alters sleep-wake architecture. (A) Experimental design for the chemogenetic re-activation of R-PSOP-responsive neurons. AAV9-hSyn-DIO-hM3D(Gq)-mCherry was injected bilaterally into the PVH of Fos2A-iCreER mice. Leaving 3 weeks for viral expression and habituation to EEG implantation, mice received 4-OHT (20mg/kg) and R-PSOP (100mg/kg) i.p.at ZT14 to induce Cre-dependent expression of the excitatory DREADD receptor explicitly in R-PSOP-responsive neurons (depicted by the red neuron) at the PVH. One week later, and after baseline day recordings (BL), animals received an i.p.injection of either saline or clozapine-N-oxide (CNO; 4mg/kg) at ZT14 in a cross-over design, followed by a recovery period (Rec). Following the completion of the recordings, an immunofluorescent validation of the injection site was performed. (B) Representative immunofluorescent image of a Fos2A-iCreER mouse expressing hM3D(Gq)-mCherry in the paraventricular hypothalamus (PVH; white dashed lines). Overlap of DAPI nuclear stain (blue) and endogenously expressed mCherry (red) suggests that both the capturing of R-PSOP-responsive neurons, and injection of hM3D(Gq) at the PVH were successful. Scale bar = 100 μm. (C) Hourly dynamics of locomotor activity on the day of injection. Blue circles and red triangles represent saline and CNO -treated groups, respectively. Gray shaded areas indicate dark phases, while the yellow shaded area represents the injection window. No statistically significant differences were observed (2-rANOVA, p <0.05; n = 5/group). (D) Hourly dynamics of wake (top), NREMS (middle), and REMS (bottom) for the respective injection day as in (C). Blue circles and red triangles represent vehicle and R-PSOP-treated groups, respectively. As before, gray shaded areas indicate dark phases and the yellow shaded area indicates the injection window. No statistically significant differences were observed (2-rANOVA, p <0.05; n = 5/group). E) Total time spent in wake, NREMS, and REMS states within the 3 hours following injection. Individual data points are shown for each animal in the saline (blue circles) or CNO (red triangles) -treated group. Box plots show median and interquartile range with individual data points. Statistically significant differences are indicated above brackets in red (paired t-tests, p<0.05; n = 5/group). (F) Heatmap summarizing changes in wake, NREMS and REMS transition probabilities (% of total state epochs) between saline and CNO, for the 3 hours following injection at ZT14 (left panel). For each transition, the starting epoch state is indicated on the left, while the ending epoch state is indicated on top. Color scale indicates magnitude and direction of changes in transition probability (yellow: increase; purple: decrease).On the right panel, individual animal transitions from the respective wake-to-NREMS and NREMS-to-wake states are shown for the saline (blue circles) or CNO (red triangles) group. Individual paired data points are connected by lines. Statistical significance is indicated in red above the bracket (paired t-test, p<0.05; n = 5/group). (G) Representative EEG spectrograms from the same mouse during the hour prior to and the first 3 hours following saline (top) or CNO (bottom) administration. Red arrowheads indicate time of injection (ZT14). Color scale represents log power. (H) Wake (left panel) and NREMS EEG (right panel) spectral power (ZT14-16; % of BL reference; see Methods) across frequencies (Hz) following saline (blue line) or CNO (red line) administration. Shaded areas represent SEM. Dark red horizontal line indicates frequencies with significant differences between treatments (Mixed effects model for ZT14-24; post-hoc Bonferroni, p<0.05; n = 5/group). (I) NREMS EEG fast delta (δ2) power (2.5-4.5Hz; normalized to d2 during BL ZT8-12; also see Methods) throughout the injection day. Blue- and red-shaded lines represent saline and CNO-treated groups, respectively. Gray shaded areas indicate dark phases, while the yellow shaded area indicates saline or CNO administration time. Statistically significant observations are shown by the dark red horizontal line (Mixed effects model for ZT14-24; post-hoc Bonferroni, p<0.05; n = 5/group). See also **Figure S5**.

## DISCUSSION

While NMUR2 has recently been implicated in various physiological processes, its specific role in sleep-wake regulation and the neural circuits mediating its effects have remained largely unexplored. In the current study, we explored the role of NMUR2 in the context of sleep and circadian rhythms in mice through the application of its only potent antagonist, R-PSOP. We provide the first comprehensive characterization of the pharmacological profile of R-PSOP in sleep homeostatic and circadian contexts. More specifically, we demonstrate dose-dependent effects of NMUR2 antagonism on LMA and sleep architecture under baseline and homeostatic challenge conditions, as well as its involvement in circadian entrainment, supporting the central role for NMUR2 in sleep regulation and circadian processes. By combining tissue clearing with neuronal tagging and pharmacology, we identify for the first time neuronal populations affected by R-PSOP throughout the diencephalon, particularly in the PVH and ARC. Our chemogenetic reactivation approach of R-PSOP-responsive PVH neurons confirms their sufficiency to drive the specific effects of R-PSOP on wake-to-NREMS transitions, NREMS amounts and NREMS EEG spectrum. The combination of our approaches represents a powerful strategy for characterizing neural targets of novel pharmacological compounds^24^ and establishes NMUR2 as a potential therapeutic target for insomnia, as well as circadian disorders marked by disrupted sleep-wake cycles.

### Physiological NMUR2 function is essential for NREM sleep and wake stability under both baseline and homeostatically challenged conditions

Delivery of NMS into the central nervous system increases arousal and wakefulness while enhancing state maintenance^8^, implicating the NMS signaling pathway in sleep-wake regulation. Our findings are complementary to these observations, suggesting that physiological NMUR2 function is essential for sleep and wake stability under both baseline and homeostatic challenge conditions. In our hands, NMUR2 antagonism primarily disrupted state stability rather than promoting a particular vigilance state, with lower doses of R-PSOP not affecting NREM sleep maintenance. Animals entered NREM sleep more frequently but failed to reach deep sleep, as evidenced by reduced d2 and spindle activity. NMUR2 function also appears to gate REMS, as evidenced by the consistent reduction in REM episodes following antagonist treatment, regardless of the circadian or homeostatic context. Notably, while effects on episode duration and number remain consistent across circadian timing, the net impact on sleep-wake amounts varied with circadian phase. We hypothesize this difference stems from directional bias in state transitions due to broader circadian influence, with wake-to-NREMS transitions predominantly affected during the active period and NREMS-to-wake transitions during the rest period. To further explore this circadian-homeostatic interaction, we examined NMUR2 antagonism under elevated homeostatic sleep pressure. Our approach revealed that R-PSOP’s effects under sleep deprivation more closely resemble those during the active period than the resting period. This suggests that the observed opposing effects may primarily reflect sleep pressure rather than circadian timing. The specific impact on faster delta oscillations, predominantly of thalamocortical origin, indicates that NMUR2 antagonism selectively affects EEG components previously associated with homeostatic challenges. Taken together, the multiple behavioral phenotypes, dose-dependent effects, and pattern of neuronal activation observed following R-PSOP administration in our study suggest that diverse structures contribute to the observed behaviors, with neuronal ensembles modulating wake-to-NREMS transitions potentially exhibiting heightened sensitivity to NMUR2 binding.

### A working model for NMUR2 antagonism

Previous studies have demonstrated NMUR2 expression throughout the hypothalamus^25^, but the specific neuronal populations mediating its behavioral effects remained unclear. Our anatomical analysis revealed that R-PSOP activated multiple structures in the mouse hypothalamus, and particularly PVH and ARC neurons, suggesting that NMUR2 acts on diverse neural components to regulate behavior. Importantly, our findings indicate that different NMUR2-mediated behaviors involve distinct neural pathways. Sleep architecture effects appear to involve SCN-NMS dependent mechanisms, likely mediated through PVH neurons, whereas LMA effects appear independent of both SCN and PVH, potentially involving lateral septum circuits^11^. These results complement recent studies implicating DMH and MPOA in NMS signaling^7^ by identifying PVH neuronal populations expressing NMUR2 as critical regulators of wake and NREMS stability. While the modulatory properties of these populations likely include inhibitory components, detailed characterization requires future investigation. Collectively, our findings suggest that hypothalamic structures including the PVH and ARC form core elements of a circuit that integrates temperature^26^, metabolic^27^, and sleep homeostatic information via NMUR2 signaling to optimize sleep and wake behavior throughout the circadian cycle.

### NMUR2 as a therapeutic target for sleep and circadian disorders

The therapeutic potential of NMUR2 as a drug target has been recognized for over two decades^12,15,28^, with agonists showing promise for obesity and metabolic disorders, while antagonists may benefit conditions including anorexia, chronic pain, osteoporosis, and stress-related disorders. Our study significantly advances our knowledge of NMUR2 function by demonstrating that R-PSOP^15^, as a potent and selective NMUR2 antagonist despite its pharmacokinetic limitations demonstrated here, effectively modulates sleep architecture and circadian entrainment *in vivo*. These findings establish NMUR2 antagonism as a viable therapeutic strategy for sleep and circadian disorders, particularly ones marked by sleep initiation difficulties. The chronopharmacological properties and neural circuits affected in our hands, together with previous observations, not only highlight the complex biology underlying NMUR2 function but also emphasize the importance of optimized dosing schedules for future NMUR2-based therapeutics. Its temporal specificity may prove particularly valuable for metabolic applications, where drug administration timing could enhance therapeutic efficacy.

Ultimately, our work positions NMUR2 as a central regulator of sleep-wake maintenance^29^ and opens new possibilities for precision medicine approaches in sleep therapeutics, where circuit-specific targeting could address the diverse manifestations of sleep and circadian disorders.

## MATERIALS & METHODS

### Stable Cell Culture and Calcium Flux Assay using Fluorescent Imaging

Chem-1 cell lines expressing the human NMUR1 (#HTS062C) and NMUR2 (#HTS098C) were bought from DiscoverX (Celle-Lévescault, France) and cultivated in DMEM, 20mM Hepes, with 10% fetal bovine serum (FBS) and 250 μg/mL geneticin. CHO cells expressing mouse NMUR1 or NMUR2 were generated by stable transfection of a plasmid containing the coding sequence. The selected stable clones were grown in F-12 K, containing 10% FBS, 1% penicillin−streptomycin, 1% L-glutamate, 300 μg/mL geneticin at 37 °C in a 10% CO_2_ incubator at 95% humidity. Cells were plated for 24 h at 50,000 cells/well in clear bottomed 96 well plates and were dye loaded for 60 min with 2 μM Fluo-4-AM (Life technologies) plus Amidoblack in assay buffer (1x HBSS (Gibco), 20mM Hepes, 2.5 mM Probenicid). The plate was loaded on a fluorometric imaging plate reader (FLIPR), compound dilution series added to the cells. After 20 min of incubation with compound, a concentration of NeuromedinS (#4048153, Bachem, Switzerland) giving 80% of the maximum signal was added to the plate and the calcium signal recorded for 5 min in order to detect antagonist activity of the test compound.

The calcium signal reduction due to the antagonist activity of the compounds was fitted to a single site competition equation with variable slope and formula Y=Bottom + (Top-Bottom)/(1+10^((LogIC50-X)*HillSlope)), where Y is the % normalized fluorescence, Bottom is the minimum Y, Top is the maximum Y, IC50 is the concentration inhibiting 50% of the agonist induced fluorescence, X is the logarithm of the concentration of the competing compound, and Hillslope the Hill coefficient. R-PSOP was tested at least 6 times in duplicate.

### Animals and housing conditions

The following lines were used: C57BL/6JRj, *Nms-iCre* (JAX #027205), *Fos^2A-iCreERT2^* (JAX #030323) and *Fos^2A-iCreERT2^* bred in-house with *Ai14*^(RCL-tdT)-D^ (JAX #007914). All experiments were performed on 10-14 week old male mice. Mice were single-housed in T3 open-top transparent type cages with *ad libitum* food and water at 12:12 LD and constant temperature (23°C) conditions. For sleep and activity experiments, behavior of mice was recorded continuously. All animals were treated according to the regulations and guidelines approved by the Veterinary Office of the Canton of Zurich.

### Surgeries

Animals received an intraperitoneal injection of Temgesic® buprenorphinium (0.1 mL/ kg) 30 minutes before surgery and were anesthetized using 1.5-2% isoflurane (Attane™ Isoflurane) dissolved in 100% oxygen with a flow rate of 0.8 L/min. Following fixation to the stereotactic system using cheek bars, the skull was calibrated and vitamin A (Hylonight®) was added to the eyes to prevent drying.

### AAV stereotactic injections

A Hamilton 7000, 0.5 μL syringe was used for all stereotactic injections. For the SCN neuronal silencing, *NMS-iCre* mice were injected at a rate of 20 nl/min with 50 nl of AAV9.hSyn1.Flex.TetLC.mCherry (UZH Vector Core, 6.4 × 10E12 viral particles/ml) at ML: +-0.18 mm, AP: −0.46 mm and DV: −5.8 mm from the bregma. For the PVH neuronal reactivation, *Fos^2A-iCreERT2^* mice were injected at a rate of 20 nl/min with 50 nl of AAV9.hSyn1.Flex.hM3D(Gq).mCherry (UZH Vector Core, 4.6 × 10E12 viral particles/ml) at ML: +-0.18 mm, AP: −0.5 mm and DV: −4.5 mm from the bregma. Following injection in each nucleus the needle was left undisturbed for another 5 minutes before being slowly retracted. The wound was stitched together with sutures (Coated VICRYL™ (polyglactin 910) sutures) and disinfected. The animals were kept on a heating mat until consciousness was recovered and provided with water containing analgesics. Post-experiment, sites of injection were histologically confirmed and unsuccessfully targeted animals were excluded from further analyses.

### EEG/EMG implantations

After animals had fully recovered for two weeks following the AAV injections, they were prepared for EEG/EMG implantation, following the same anesthetic and analgesic process as above. Briefly, gold-plated screws were placed at ML: −1.0 mm, AP: 1.5 mm from the bregma for the frontal electrode, at ML: 1.7 mm, AP: −2.5 mm for the parietal electrode, and at ML: 0.0 mm, AP: −6.0 mm for the referential electrode. EMG wires were inserted into the trapezius (neck) muscles and the EEG wires were wrapped around the screws of the corresponding positions. The screws together with the connecting wires were finally cemented to the skull, and mice were left to recover for an additional week prior to experimentation.

### EEG recordings and analysis

The EEG and EMG signals were amplified (amplification factor, _2000), filtered (high pass filter: –3 dB at 0.016 Hz; low pass filter: –3 dB at 40 Hz) sampled with 512Hz, digitally filtered [EEG: low pass finite impulse response (FIR) filter, 0.1-25Hz; EMG: bandpass FIR filter, 10-30Hz], and stored with a final resolution of 128 Hz. Sleep was scored by computer assisted staging (SPINDLE^30^) followed by visual inspection, unless stated differently. Data analysis was performed with custom-made MATLAB scripts (MathWorks). EEG spectrograms were visualised with a custom-made script built on MNE-Python and YASA 0.6.5.

### LMA recordings and analysis

Data on LMA was collected with infrared sensors placed on top of the cage, recorded and analyzed with ClockLab4 Wireless software (Actimetrics). For LMA graphs, activity was sorted into 60 min bins and the average behavior over the 24-hour baseline day for each mouse was used as normalization reference to allow comparisons between animals.

### PFA and aCSF perfusion

For PFA perfusions, fresh 4% PFA was prepared by diluting a 20% PFA stock solution in phosphate buffer (PB). Mice were anesthetized using 2-3% isoflurane and euthanized using 200 mL Pentobarbital Natrium. The bodies were fixed on a board with pins, and the heart was revealed by opening the chest area and removing the ribcage. A small incision was cut in the right ventricle, and first, 10 mL of PBS were run through the blood circulation, followed by 10 mL of 4% PFA. The brains were extracted and fixed in 10 mL 4% PFA for 2 h to overnight. Then the brains were stored in 30% sucrose diluted in 0.5 M PB for at least three days until further use. For aCSF perfusions, fresh aCSF solution was prepared by diluting a 10x aCSF stock solution in distilled water with the addition of glucose, CaCl_2_ and MgCl_2_ to a final concentration of 25 mM glucose, 2.5 mM CaCl_2_ and 2 mM MgCl_2_. The aSCF solution was cooled on ice and aerated with carbogen gas for 30 mins. The pH was adjusted to 7.4 and until the end of the perfusion the solution was kept on ice aerated with carbogen gas. Animals were perfused similarly as described in the PFA perfusion. Brains were extracted, whereas the cerebellum and the olfactory bulb were removed and the remaining brain was cut in half and post-fixed in freshly prepared 4% PFA for 2 hours. Brains were rinsed with cold PBS before storage in 30% sucrose.

### Immunofluorescent stainings

Brains were fixed on a cryostat base at −21°C using the frozen section medium Neg-50^TM^. After calibration, brain slices were taken with a 40 µm thickness and kept in an anti-freezing solution until further use. At the day of staining, brain slices were washed 2 × 10 mins in 2 mL Tris-Triton 0.05% at RT on a shaker at 100 rpm and subsequently blocked in 400 µL NDS or NGS 5% (depending on whether the host of the secondary antibody was donkey or goat) in Tris-Triton 0.05% for one hour at RT on a shaker at 75 rpm. The slices were then incubated in 400 µL primary antibody solution (anti-mCherry (Sicgen AB0081-200) diluted 1:1000; anti-cFOS (Sysy Cat. No. 226 017) diluted 1:1000, overnight in a moist chamber at 4°C on a shaker at 75 rpm and afterwards washed with 2 mL Tris-Triton 0.05% as before. The following day they were incubated in 400 µL secondary antibody solution (anti-goat Alexa Fluor™ 594 diluted 1:1000; anti-rat Alexa Fluor™ 488 diluted 1:1000, anti-rabbit Alexa Fluor™ 594 diluted 1:1000) for one hour at RT on a shaker at 75 rpm/min and in 2 mL 1:10’000 DAPI in Tris-Triton 0.05% for 10 mins at RT on a shaker at 100 rpm/min, prior to two final washing steps. The brain slices were then mounted on gelatin-coated glass slides, mounting media (Dako Fluorescence Mounting Medium) was applied to the slides, a glass cover slip was placed on each slide, and sealed with clear nail polish.

### Mouse intraperitoneal single dose pharmacokinetic with R-PSOP and selectivity screening

An exploratory single-dose pharmacokinetic study in three C57BL/6J male mice was conducted with R-PSOP following intraperitoneal administration at the dose of 100mg/kg. The compound was formulated with 10% hydroxypropyl cyclodextrin in 0.7% NaCl. Plasmasamples were collected at 6 time points: 0, 0.25, 0.5, 1, 2, 5 hours and CSF was collected 1 hour after administration. The compound was detected using mass spectrometry and pharmacokinetic parameters were derived from the plasma concentrations versus time profile as well as the CSF concentration.

Using *in vitro* plasma free fraction (fup = 0.36), unbound brain/plasma partition coefficient Kp,uu= 0.15 and *in vitro* NMUR2 binding data (mouse NMUR2 Ki = 197 nM, N= 6), the following Emax equation was used to calculate brain receptor occupancy at Cmax (assuming competitive inhibition):

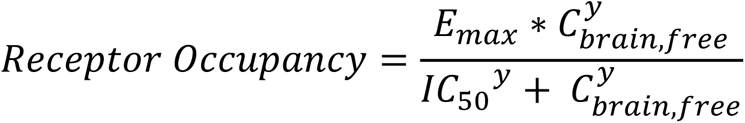

where Emax is the maximum binding, assumed to be 100%, the hill coefficient y is assumed to be 1, C_brain,free_ = concentration unbound, free in the brain, and IC50 represents *in vitro* binding to the receptor.

Selectivity screening was performed at Eurofins Cerep (Celle l’Evescault, France) on a panel of 31 human and 5 rat receptors as well as in 13 human and 1 bovine enzyme assays.

### 4-OHT preparation and administration

For 4-Hydroxytamoxifen (4-OHT) preparation we followed a protocol developed by the Deisseroth lab (https://web.stanford.edu/group/dlab/optogenetics/sequence_info.html). Briefly, 4-OHT powder was diluted in 250ul DMSO (Sigma H6278). Subsequently, the diluted 4-OHT was mixed with 4.75 ml of separately prepared 25% Tween80 diluted in saline, and mixed by vigorous vortexing. The resulting clear solution was used within the next 1 hour for intraperitoneal (i.p.) injections of 250 μl volume per animal to deliver a final concentration of 20mg/kg 4-OHT. For the reactivation experiment, tagging of NMUR2 responsive neurons was achieved via simultaneous i.p. injection of 20mg/kg 4-OHT in saline and 100mg/kg R-PSOP at ZT14 in *Fos^2A-iCreERT2^* mice injected with hM3D(Gq) -expressing AAV at the PVH. Following seven days for Gq expression in tagged neurons, mice were fully habituated and ready for chemogenetic experimentation. For the 3D brain clearing experiment, tagging of NMUR2 responsive neurons was achieved via the same protocol, this time in *Fos^2A-iCreERT2^*;*Ai14* mice. Following seven days for the expression of tdTomato in captured neurons, mice were terminated via pentobarbital administration, perfused, and their brains were extracted and underwent the CUBIC tissue-clearing process.

### CUBIC-HV tissue clearing

The CUBIC whole brain clearing pipeline was followed as previously described. Briefly, whole brains were immersed in 4% paraformaldehyde (PFA) for 24h, then delipidated and decolored for 5 days in CUBIC-L (CUBIC-HV, TCI chemicals). Subsequently, their cellular nuclei were stained with DAPI for another 5 days, and the refracting index (RI; 1.520) of the transparent brain was calibrated in CUBIC-R+ solution (CUBIC-HV, TCI chemicals) for three days. Following the end of incubations and in-between washes, the cleared brains were stored for two days in mineral oil (Mounting Solution (RI 1.520), TCI chemicals) and subsequently imaged with axially-scanned light-sheet imaging.

### Image acquisition with mesoscale selective plane imaging microscopy (mesoSPIM)

The mesoscale selective plane illumination microscopy instrument (mesoSPIM, https://mesospim.org/) was used for axially-scanned light-sheet imaging of the cleared brains. Transparent brain images were recorded at 3.2x magnification with an Olympus MVX-10 macroscope with a MVPLAPO objective combined with a Hamamatsu ORCA-FLASH 4.0 V3 camera, at a voxel size of 2.03 × 2.03 × 5 μm3 (X × Y × Z). The laser/filter combinations for imaging were as follows: for registration of autofluorescence and light-scattering, a 405nm (100mW) excitation laser at 100% and no emission filter. For the tdTomato signal, a 561nm (100mW) excitation laser at 50% and a quadruple bandpass (BP444/27; BP523/22; BP594/20; BP704/46) emission filter. Brains were positioned such that the ventral portion was the first plane acquired. Briefly, each whole brain was imaged in 16 tiles per channel (eight tiles per illumination; right or left). The acquired data occupied 100GB per brain per channel on average.

### 3D image processing and visualization of cleared brains

Following acquisition, images were stitched and fused separately for each channel using the FIJI plugin BigStitcher^31^, according to the “MesoSPIM stitching using BigStitcher” guidelines (https://zmb.dozuki.com/Guide/MesoSPIM+Stitching+using+BigStitcher/283). Finally, stitched brain images were stored in .tiff format for visualization with napari (Python)^32^. Parameters in napari were as follows: opacity = 1; blending = translucent_no_depth; auto-contrast = continuous; gamma = 0.69; colormap = twilight; interpolation = linear. For the axial, sagittal and coronal views of the brain shown in 3D, slice 25/103, 77/159 and 71/191 were used.

## QUANTIFICATION AND STATISTICAL ANALYSIS

### Statistical analysis

All statistical analyses, including two-way RM ANOVAs, paired t-tests and Tukey’s tests were performed with PRISM 9 (GraphPad). All tests were applied as described in figure legends. Statistical significance was defined as p < 0.05, where p values are not stated.

#### Lead contact

Further information and requests for resources and reagents should be directed to and will be fulfilled by the lead contact, Dr. Konstantinos Kompotis and Dr. Christophe Grundschober.

## ACKNOWLEDGMENTS

We would like to thank Dr Alex Rosi-Andersen for his hands-on assistance and Dr Shiva Tyagarajan for his intellectual contributions on the underlying circuitry, as well as to Stefan Berchtold for his support on the compound preparation. The study was supported by a Velux Stiftung grant to SAB and KK and a Research grant from Hoffmann-La Roche to SAB. This study is dedicated to Prof. Steven A Brown.

## AUTHOR CONTRIBUTIONS

*Conceptualization:* K.K., S.A.B., C.G, and R.L.R.; *methodology:* K.K., S.A., M.S, and M.B.; *investigation:* K.K, S.A., M.S., S.G. and B.D.V.G.; *formal analyses*: K.K, M.S, B.D.V.G. and S.G.; *visualization*: K.K and M.S.; *writing—original draft:* K.K. and S.A.; *writing—review & editing:* M.S., R.M.R.S., C.G. and R.L.R.; *funding acquisition:* K.K., S.A.B. and C.G.; compounds preparation, evaluation of properties and pharmacokinetics for the proper use of R-PSOP: RMRS., BDVG.; *resources:* K.K, S.A.B., R.M.R.S. and R.L.R.; *supervision:* K.K., S.A.B., C.G., and R.L.R. All authors approved the final version of the manuscript.

## DECLARATION OF GENERATIVE AI AND AI-ASSISTED TECHNOLOGIES

Nothing to declare.

## SUPPLEMENTAL INFORMATION

**Document S1. Table S1**

**Document S2. Figures S1–S5** sample

---

**Figure S1.**
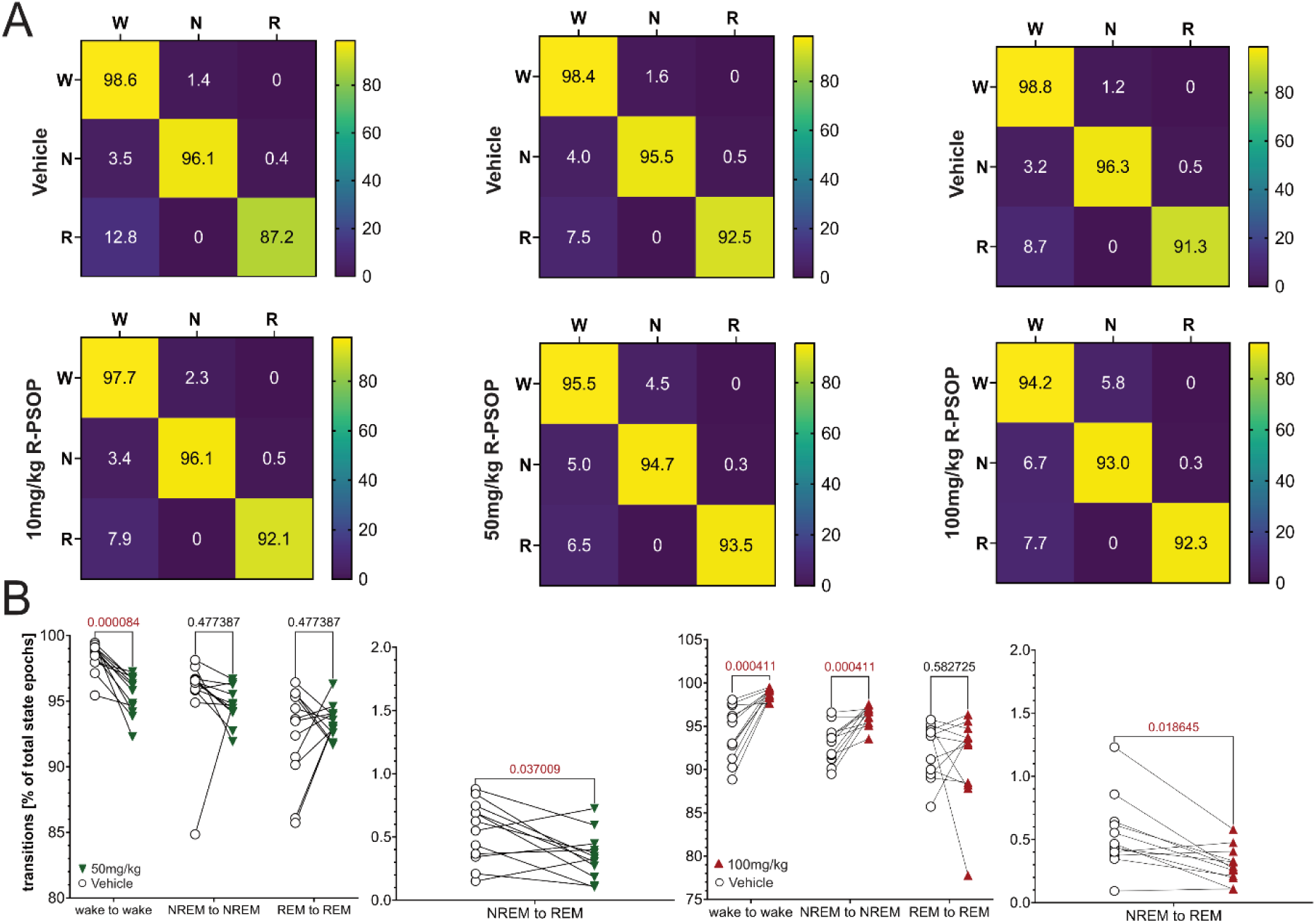
R-PSOP affects locomotor activity and sleep architecture in a dose-dependent manner. Related to Figure 2. **A)** Heatmaps showing wake, NREMS and REMS transition probabilities (% of total state epochs) for vehicle and R-PSOP at 10mg/kg (left), 50mg/kg (middle), and 100mg/kg (right), during the 4-hour period following injection. For each transition, the starting epoch state is indicated on the left, while the ending epoch state is indicated on top. Color scale indicates magnitude and direction of changes in transition probability (blue: low; yellow: high). **B)** Individual animal transitions for the same vigilance state (left) and NREMS-to-REMS (right) transitions for the 50mg/kg and 100mg/kg and respective vehicle groups. Corresponding p-values for significant differences are shown above brackets in red (paired t-tests, p<0.05; n = 12).

**Figure S2.**
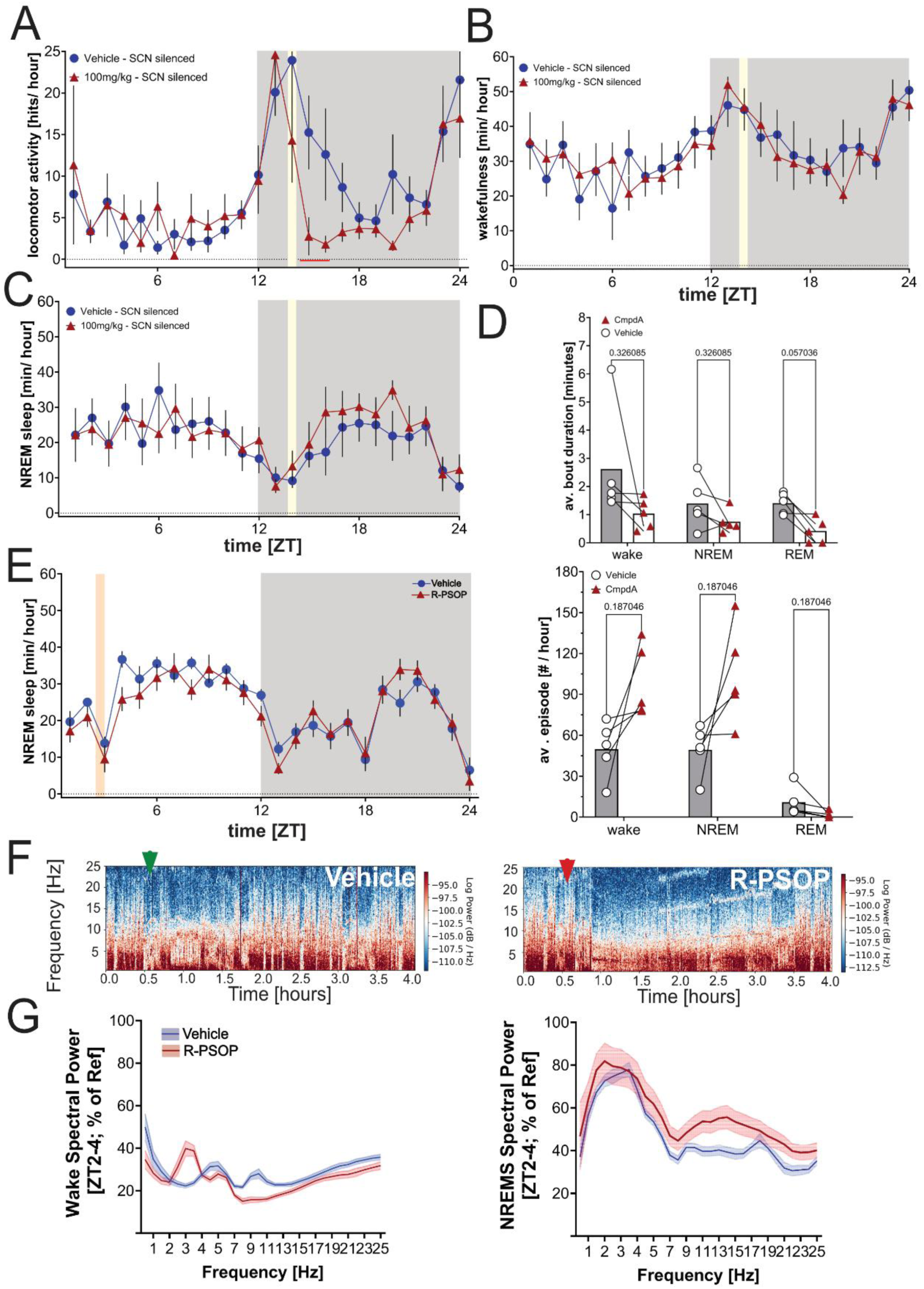
Effects of NMUR2 antagonism during the resting period on circadian entrainment and sleep-wake behavior. Related to Figure 4. Time course of **A)** locomotor activity, **B)** wakefulness and **C)** NREM sleep over 24 hours in mice treated at ZT14 with vehicle (blue circles) and 100mg/kg (red ascending triangles) of R-PSOP in SCN -silenced mice (n=5). Data points represent mean ± SEM. Gray shaded areas indicate dark phase; yellow vertical lines indicate time of intraperitoneal injection. Horizontal red bar indicates significant differences between vehicle and R-PSOP-treated group (2-rANOVA, post-hoc Tukey tests, p < 0.05; n = 12). **D)** (Top panel) Average episode duration (top panel) and number (bottom panel) of wake, NREMS, and REMS in the 4 hours following administration of vehicle or R-PSOP in SCN-silenced mice. Individual data points are shown as empty circles and red ascending triangles, respectively. Individual paired data points are connected by lines. Bar graphs represent mean ± SEM. Statistically significant differences are indicated above brackets in red (paired t-tests, p<0.05; n = 5). **E)** Time course of NREM sleep over 24 hours in mice treated at ZT2 with vehicle (blue circles) or 100mg/kg R-PSOP (red ascending triangles). Data points represent mean ± SEM. Gray shaded areas indicate dark phase; yellow vertical line indicate time of intraperitoneal injection. No significant differences between vehicle and R-PSOP-treated group were observed (2-rANOVA, post-hoc Tukey tests, p > 0.05; n = 5). **F)** Representative raw EEG spectrograms depicting the hour prior to and the first 3 hours following administration of Vehicle (left panel) or 100mg/kg R-PSOP at ZT2 (right panel). Red arrowheads indicate time of injection. Color scale represents log power. **G)** Wake (left panel) and NREMS EEG (right panel) spectral power (ZT2-4; % of BL reference; see Methods) across frequencies (0.5-25Hz) following saline (blue line) or CNO (red line) administration. Shaded areas represent SEM. Dark red horizontal line indicates frequencies with significant differences between treatments (Mixed effects model for ZT2-4; post-hoc Bonferroni, p>0.05; n = 5/group).

**Figure S3.**
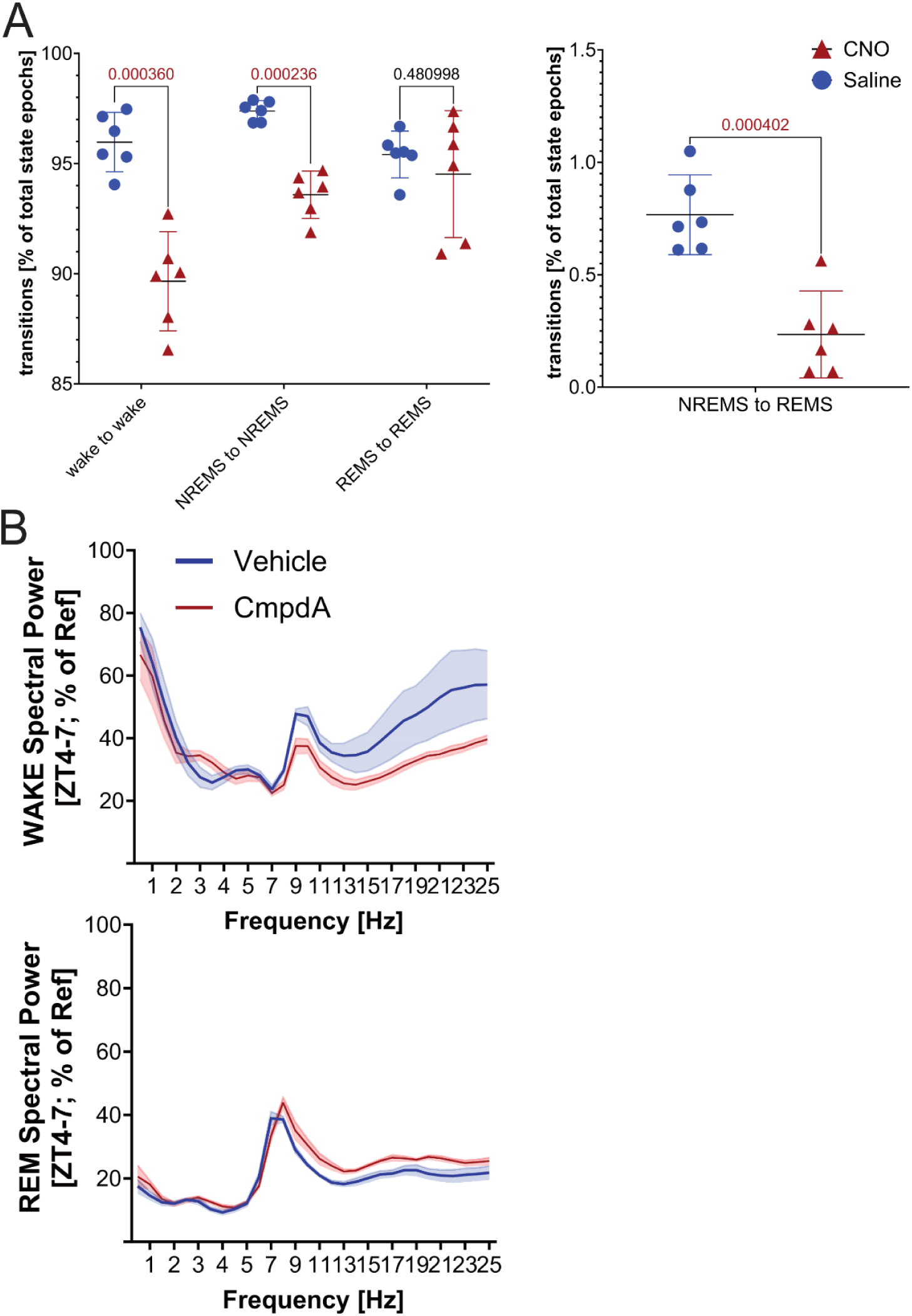
Administration of R-PSOP alters homeostatic response to forced wakefulness. Related to Figure 5. **A)** Individual animal transitions to the same state (left) and from NREMS-to-REMS (right) in the 3 hours following SD. Corresponding p-values for significant differences are shown above brackets in red (t-tests, p<0.05; n = 6/group). **B)** Wake (top panel) and REMS EEG (bottom panel) spectral power (ZT4-7; % of BL reference; see Methods) across frequencies (Hz) following saline (blue line) or CNO (red line) administration. Shaded areas represent SEM. No significant differences between treatments were observed (Mixed effects model for ZT4-7; post-hoc Bonferroni, p>0.05; n = 6/group).

**Figure S4.**
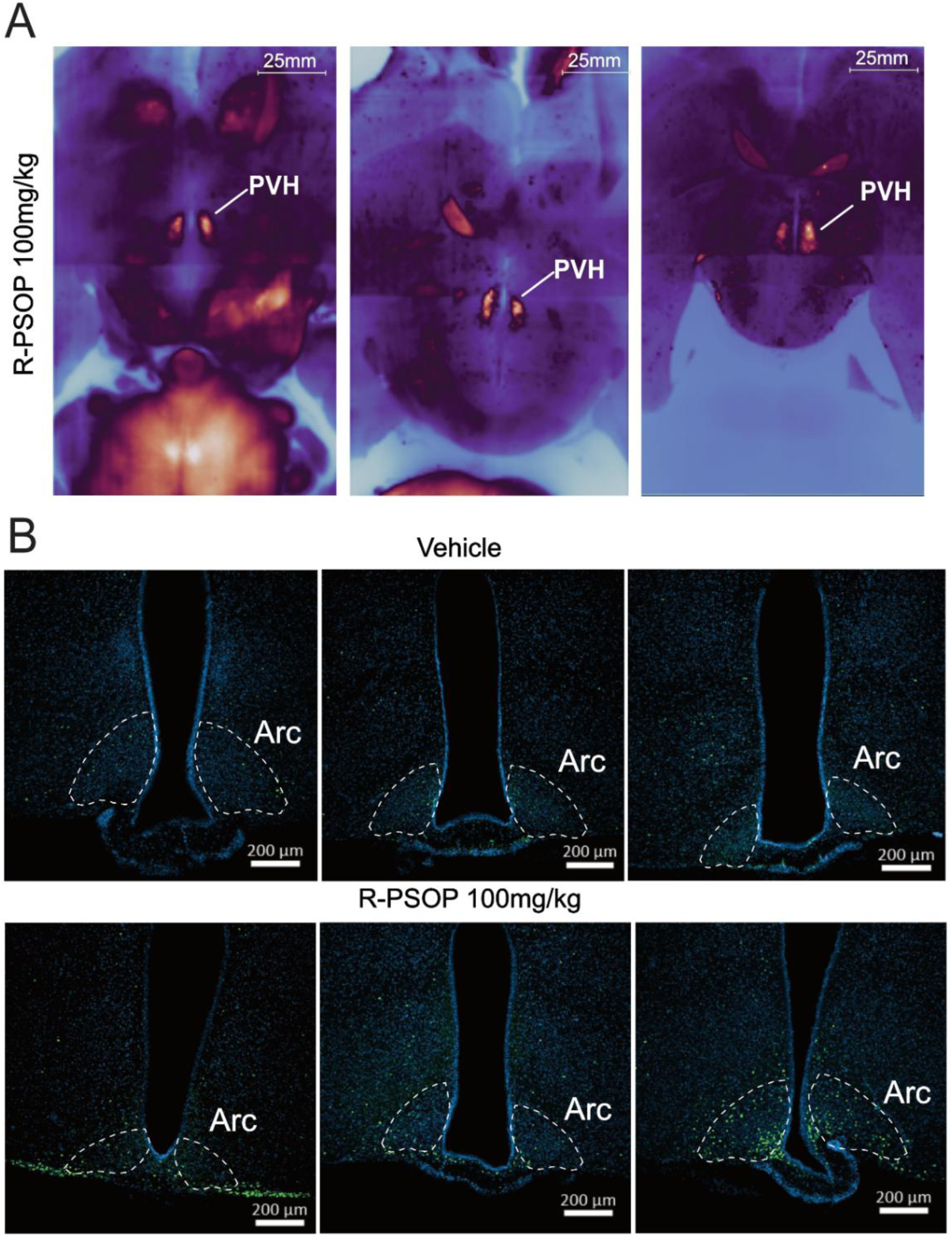
R-PSOP activates paraventricular hypothalamic and arcuate nucleus neurons. Related to Figure 6. **A)** Three-dimensional visualization of R-PSOP-activated neurons throughout the brain. Axial views for 3 individual animals show labeled neurons with particular enrichment in the paraventricular hypothalamus (PVH). Yellow/orange dots represent individual activated neurons; blue background shows brain autofluorescence for anatomical reference. Scale bar = 25 mm. Napari python package was used for visualization (also see Methods for specific parameters). **B)** Representative immunofluorescent images of the arcuate nucleus (Arc; white dashed lines) 1.5 hours after injection at ZT14 with either vehicle (top panels) or R-PSOP (100mg/kg, bottom panels). DAPI nuclear stain (blue) and cFos immunoreactivity (green) show increased neuronal activation in R-PSOP-treated brain. In each group, three individual animals are presented. Scale bar = 200 μm.

**Figure S5.**
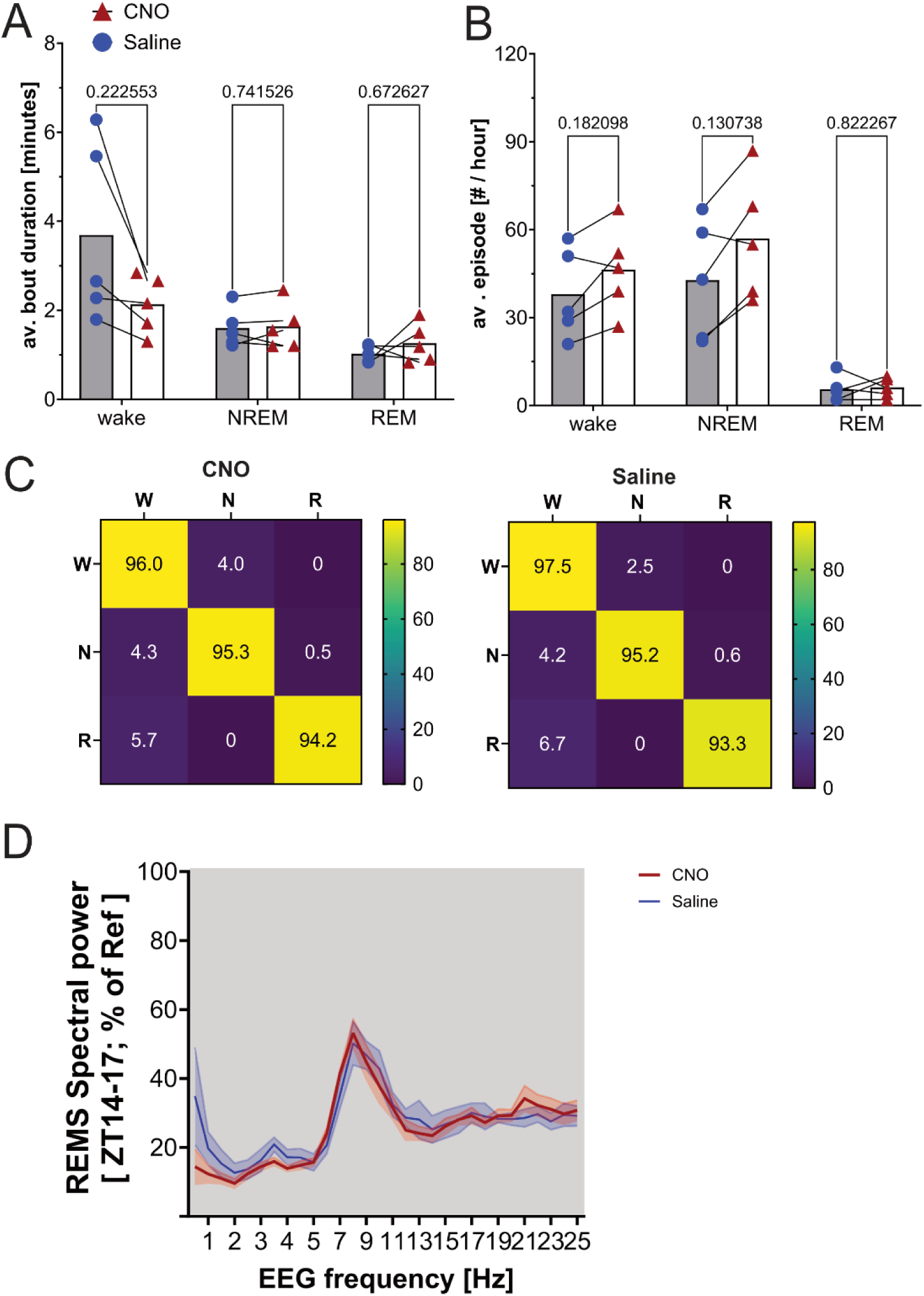
Chemogenetic reactivation of R-PSOP-responsive does not alter sleep-wake architecture. Relative to Figure 7. Average episode **A)** duration and **B)** number of wake, NREMS, and REMS in the 3 hours following administration of vehicle or 100mg/kg R-PSOP. Individual data points are shown as blue circles (Vehicle) or red ascending triangles (100mg/kg). Individual paired data points are connected by lines. Bar graphs represent mean ± SEM. No statistically significant were observed (paired t-tests, p>0.05; n = 5). **C)** Heatmap summarizing changes in wake, NREMS and REMS transition probabilities (% of total state epochs) between CNO (left) and saline (right), for the 3 hours following injection at ZT14. For each transition, the starting epoch state is indicated on the left, while the ending epoch state is indicated on top. Color scale indicates magnitude and direction of changes in transition probability (yellow: high; purple: low). **D)** REMS EEG spectral power (ZT14-17; % of BL reference; see Methods) across frequencies (0.5-25Hz) following saline (blue line) or CNO (red line) administration. Shaded areas represent SEM. Dark red horizontal line indicates frequencies with significant differences between treatments (Mixed effects model for ZT14-17; post-hoc Bonferroni, p>0.05; n = 5/group).

**Table S1:**
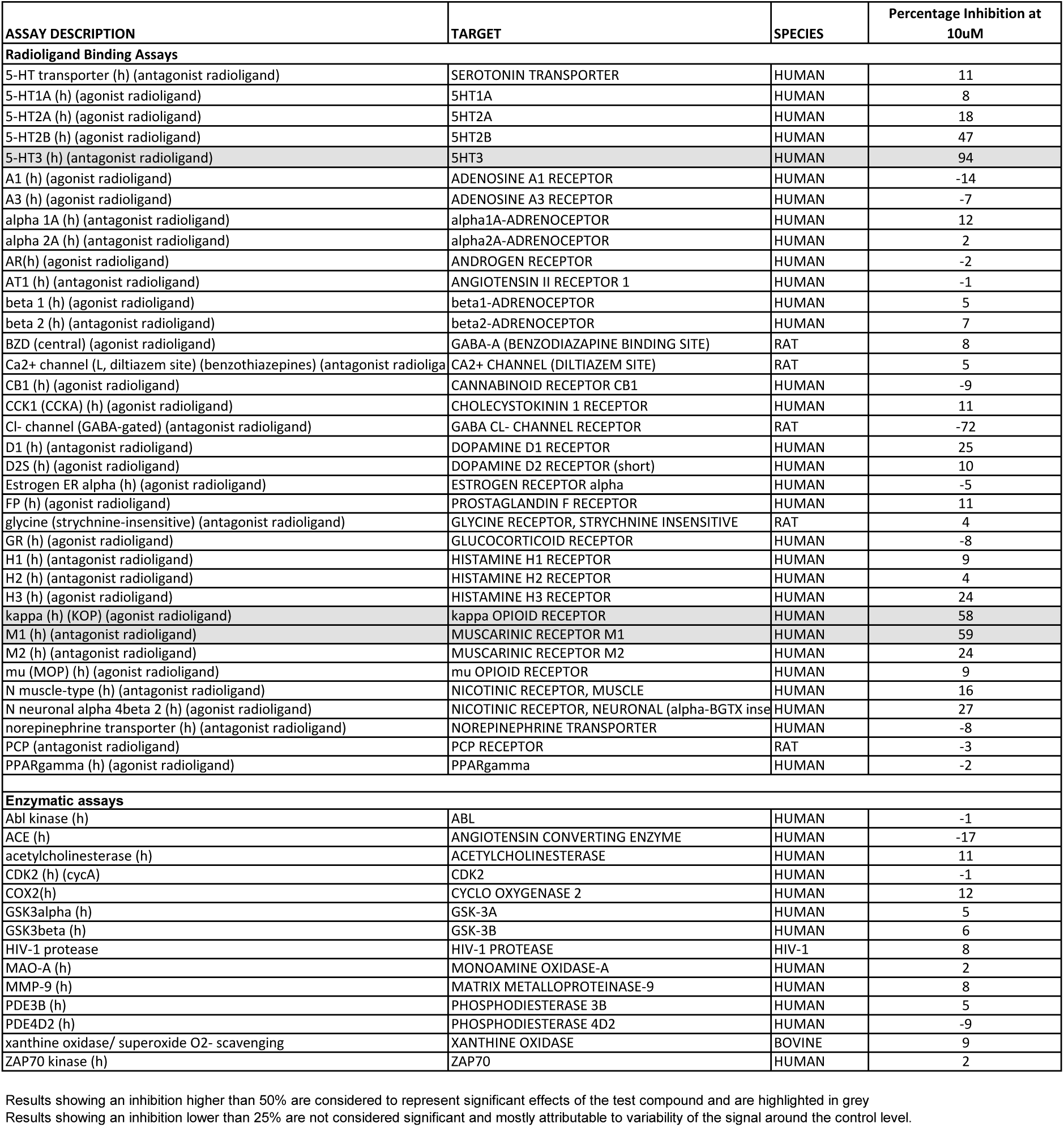
Selectivity testing of R-PSOP on human/rat receptor transporters and enzymes at 10 uM, Related to Figure 1.

